# UBXN1 maintains ER proteostasis and represses UPR activation by modulating translation independently of the p97 ATPase

**DOI:** 10.1101/2022.12.16.520763

**Authors:** Brittany A. Ahlstedt, Rakesh Ganji, Sirisha Mukkavalli, Joao A. Paulo, Steve P. Gygi, Malavika Raman

## Abstract

Endoplasmic reticulum (ER) protein homeostasis (proteostasis) is essential to facilitate proper folding and maturation of proteins in the secretory pathway. Loss of ER proteostasis due to cell stress or mutations in ER proteins can lead to the accumulation of misfolded or aberrant proteins in the ER and triggers the unfolded protein response (UPR). In this study we find that the p97 adaptor UBXN1 is an important negative regulator of the UPR. Loss of UBXN1 significantly sensitizes cells to ER stress and activates canonical UPR signaling pathways. This in turn leads to widespread upregulation of the ER stress transcriptional program. Using comparative, quantitative proteomics we show that deletion of UBXN1 results in a significant enrichment of proteins belonging to ER-quality control processes including those involved in protein folding and import. Notably, we find that loss of UBXN1 does not perturb p97-dependent ER associated degradation (ERAD). Our studies indicate that loss of UBXN1 increases translation in both resting and ER-stressed cells. Surprisingly, this process is independent of p97 function. Taken together, our studies have identified a new role for UBXN1 in repressing translation and maintaining ER proteostasis in a p97 independent manner.

## Introduction

The ER is a major site for protein maturation and oversees the folding and secretion of one-third of the cellular proteome. A tight equilibrium exists between the ER protein folding capacity and protein load to maintain ER proteostasis. Alterations in this equilibrium due to various physiological conditions such as disruption of ER-calcium homeostasis, mutations that impair protein folding, or cellular aging can result in protein misfolding^1^. ER stress is elicited by protein misfolding and aggregation and can become pathological if not alleviated. Indeed, protein misfolding and aggregation are hallmarks of many age-associated neurodegenerative disorders. Numerous studies indicate that in neurodegenerative disorders such as Parkinson’s Disease (PD) and amyotrophic lateral sclerosis (ALS), age associated protein aggregation in the ER is causally linked to cell death^2–5^. Protein aggregates can inappropriately interact with ER chaperones, disrupting ER morphology and vesicular trafficking to the Golgi apparatus^6–8^. In the context of these age-related disorders, it is known that a decline in the efficiency and capacity of protein quality control systems diminishes ER-protein homeostasis. However, it is currently unclear how decline in specific protein quality control mechanisms impacts cell health. Recent studies suggest that aging cells rewire chaperone subnetworks and find that proteasome and autophagy capacities are altered^9–11^. Thus, a thorough understanding of protein quality control mechanisms is warranted to comprehend how their decline triggers disease onset.

Distinct ER-quality control mechanisms sense ER stress. These include the unfolded protein response (UPR) and ER-associated degradation (ERAD) which function to improve the folding capacity of the ER, and clear misfolded proteins from the ER through the ubiquitin-proteasome system respectively. The UPR is an adaptive ER-quality control pathway comprised of three, parallel, ER-resident transmembrane stress sensors that monitor the misfolded protein burden within the ER. These sensors include activating transcription factor 6 (ATF6), inositol requiring enzyme 1 (IRE1), and protein kinase R (PKR)-like ER kinase (PERK). Signaling pathways downstream of each sensor ultimately activates a distinct transcription factor that induces the expression of a suite of UPR-specific target genes, encoding chaperones and protein degradation machinery^12^. The prevailing theory suggests that the ER sensors are held in an inactive state when their luminal domains are bound to the abundant ER chaperone, binding immunoglobulin protein (BiP) (also known as glucose-regulated protein 78 kDa, GRP78)^13^. When misfolded proteins accumulate, BiP is titrated away from the sensors, triggering their activation^14, 15^. Under these conditions, ATF6 traffics from the ER to the Golgi apparatus where site-1 and site-2 proteases release the cytosolic domain of ATF6 to generate an active transcription factor. In contrast, PERK dimerizes upon activation and phosphorylates eukaryotic initiation factor 2α (eIF2α), diminishing the translation initiation of most proteins^16^. In response to eIF2α phosphorylation, the translation of activating transcription factor 4 (ATF4) is selectively induced in response to reinitiation of ribosomes at the upstream *atf4* coding region. Lastly, IRE1 forms oligomers and mediates the unconventional mRNA splicing of an intron from the x-box binding protein-1 (*xbp1*) mRNA to generate the active transcription factor xbp1s^17, 18^. Additionally, IRE1 degrades ER-targeted mRNAs in a process termed regulated IRE1-dependent decay (RIDD)^19^. The rapid degradation of mRNAs through RIDD functions to alleviate ER stress by preventing the continued influx of new polypeptides into the ER^20, 21^. If these mechanisms fail to alleviate ER stress, PERK and IRE1 signaling will induce apoptosis through expression of the pro-apoptotic transcription factor C/EBP homologous protein (CHOP) and JNK signaling respectively^22–24^.

The UPR works in tandem with ERAD which purges terminally misfolded clients from the ER by the ubiquitin-proteasome system. During ERAD, misfolded ER proteins are recognized and handed off to ER membrane resident E3 ligases to initiate ubiquitin-dependent degradation by cytosolic proteasomes. The AAA+ ATPase p97 (also known as valosin containing protein, VCP) is a critical intermediate in this process that recognizes ubiquitylated proteins and retro-translocates them from the ER to the cytosol prior to proteasomal degradation. The p97 homo-hexamer utilizes ATP hydrolysis to unfold substrates^25–27^ and participates in an array of cellular pathways from the cell cycle to autophagy^28, 29^. p97 interacts with a variety of dedicated adaptor proteins that facilitate substrate specificity and targeting of p97. The largest family of p97 adaptors contain a ubiquitin regulatory X (UBX) domain that enables binding to p97^30^. Many of the UBX domain containing adaptors also harbor a ubiquitin associated domain (UBA) that associates with polyubiquitin chains on client proteins. There are ∼40 adaptors, though the functions and substrates of most remain poorly characterized. Studies have shown that p97 mutation can disrupt p97-adaptor binding which has been causally linked to multisystem proteinopathy 1 (MSP-1), a degenerative disorder wherein individuals present with inclusion body myopathy, Paget’s disease of the bone and/or frontotemporal dementia^31–35^. More recently, p97 mutation has been linked to the development of familial ALS (fALS), PD, and Charcot-Marie-Tooth disease type IIB (CMTIIB)^36–39^. A full understanding of adaptor function is required to better understand how their dysfunction disrupts protein quality control pathways to contribute to disease pathogenesis.

In a previous study, we reported that the p97 adaptor UBX domain protein 1 (UBXN1) associated with the multifunctional BCL2-associated athanogene 6 (BAG6) chaperone. BAG6 facilitates the insertion of tail anchored proteins into the ER^40–45^. When clients fail to insert (due to mutations, saturation of insertion machinery at the ER, or ER stress), BAG6 recruits the E3 ligase RNF126 for ubiquitylation and degradation of the substrate^44^. We showed that UBXN1 was important for the recruitment of p97 to the BAG6 triage complex to facilitate extraction and degradation of non-inserted substrates. In addition, BAG6 has also been reported to function in ERAD in concert with p97 by acting as a chaperone ‘holdase’ for unfolded p97 clients^46–48^. However, our studies indicated that UBXN1 is not a ERAD adaptor for p97^40^. Here, we have identified a novel role for UBXN1, as a regulator or ER proteostasis. In this study, we demonstrate that cells depleted of UBXN1 have elevated UPR activation both in unperturbed and ER-stressed cells. Using quantitative proteomics, we find a significant increase in the abundance of ER proteins in UBXN1 deleted cells. Increased steady state protein levels are not due to the participation of UBXN1 in p97-dependent ERAD. We further demonstrate that loss of UBXN1 increases protein translation in a p97-independent manner and this may be the cause of ER stress. Our studies have identified a new regulator of protein translation and ER proteostasis.

## Results

### UBXN1 depletion induces activation of the unfolded protein response

Based on our prior study on p97-UBXN1 regulation of BAG6-associated protein triage, we presumed that one consequence of UBXN1 loss may be imbalance in the ER proteome and subsequent ER stress. Indeed, loss of p97 or BAG6 function can cause the accumulation of misfolded proteins in the ER via multiple mechanisms to activate the UPR. Silencing BAG6 with siRNA or inhibiting p97 with the ATP-competitive inhibitor CB-5083 caused robust activation of the UPR as expected by immunoblotting for the ER chaperone BiP and transcription factor ATF4 (Supplementary Figure 1 A-E). To assess if UBXN1 was also involved in ER stress responses, we systematically assessed UPR activation in UBXN1 knock-out (KO) HeLa Flp-In T-REx (HFT) cell lines generated by CRISPR-Cas9 gene editing. Wildtype and UBXN1 KO cells were treated with the reducing agent dithiothreitol (DTT), and lysates were probed for the ER chaperone BiP, the transcription factor ATF4, and phosphorylation of eIF2α (Figure 1A and B). In wildtype cells, a time-dependent increase in these proteins was observed, however a significant increase was apparent at earlier time points in UBXN1 KO cells. Notably, these markers were already upregulated in untreated UBXN1 KO cells (Figure 1A, compare lane 5 and 1). We next monitored the levels of cleaved ATF6 in wildtype and UBXN1 KO cells treated with DTT. Cells were co-treated with the proteasome inhibitor Bortezomib (BTZ) to prevent turnover of and facilitate visualization of the N-terminus of ATF6^49^. Compared to control cells, there was a modest but significant increase in cleaved ATF6 in UBXN1 KO cells in response to ER stress (Figure 1C and D). Under ER stress conditions, IRE1*α* is activated by dynamic clustering in the ER membrane^50, 51^. IRE1*α* monomers cluster into higher order oligomers that can be imaged and quantified by immunofluorescence. We generated a stable cell line using a previously published doxycycline-inducible GFP-tagged IRE1*α* reporter to observe IRE1*α* clustering^51^. Utilizing this cell line, IRE1 clustering into oligomers was quantified by quantifying the number of GFP puncta. UBXN1 was depleted with two independent siRNAs and cells were imaged for GFP-foci (Figure 1E). We found that 40% of cells depleted of UBXN1 contained GFP-IRE1*α* foci compared to the 10% observed in cells transfected with a control siRNA (Figure 1F). Furthermore, there were significantly more GFP-foci per cell in UBXN1-depleted cells compared to control (Figure 1F). Additionally, we measured IRE1 activation via downstream *xbp1s* expression in UBXN1 KO cells by quantitative real-time PCR using a primer set that spans the splice site removed by IRE1*α*^52^. We detected 19-fold greater *xbp1s* expression in UBXN1 KO cells compared to control in response to stress, while total *xbp1* levels remained unchanged (Figure 1G). A similar phenotype was observed in UBXN1 KO HEK-293T cells, (Supplementary Figure 1F). Notably, the repression of ER stress by UBXN1 is not due to a role for UBXN1 as a p97 adaptor for ERAD^40, 53^. Using a panel of GFP-tagged ERAD reporters we have previously shown that p97 and its adaptor UBXD8 but not UBXN1 is required for the turnover of multiple ERAD substrates^40^.

**Figure 1.**
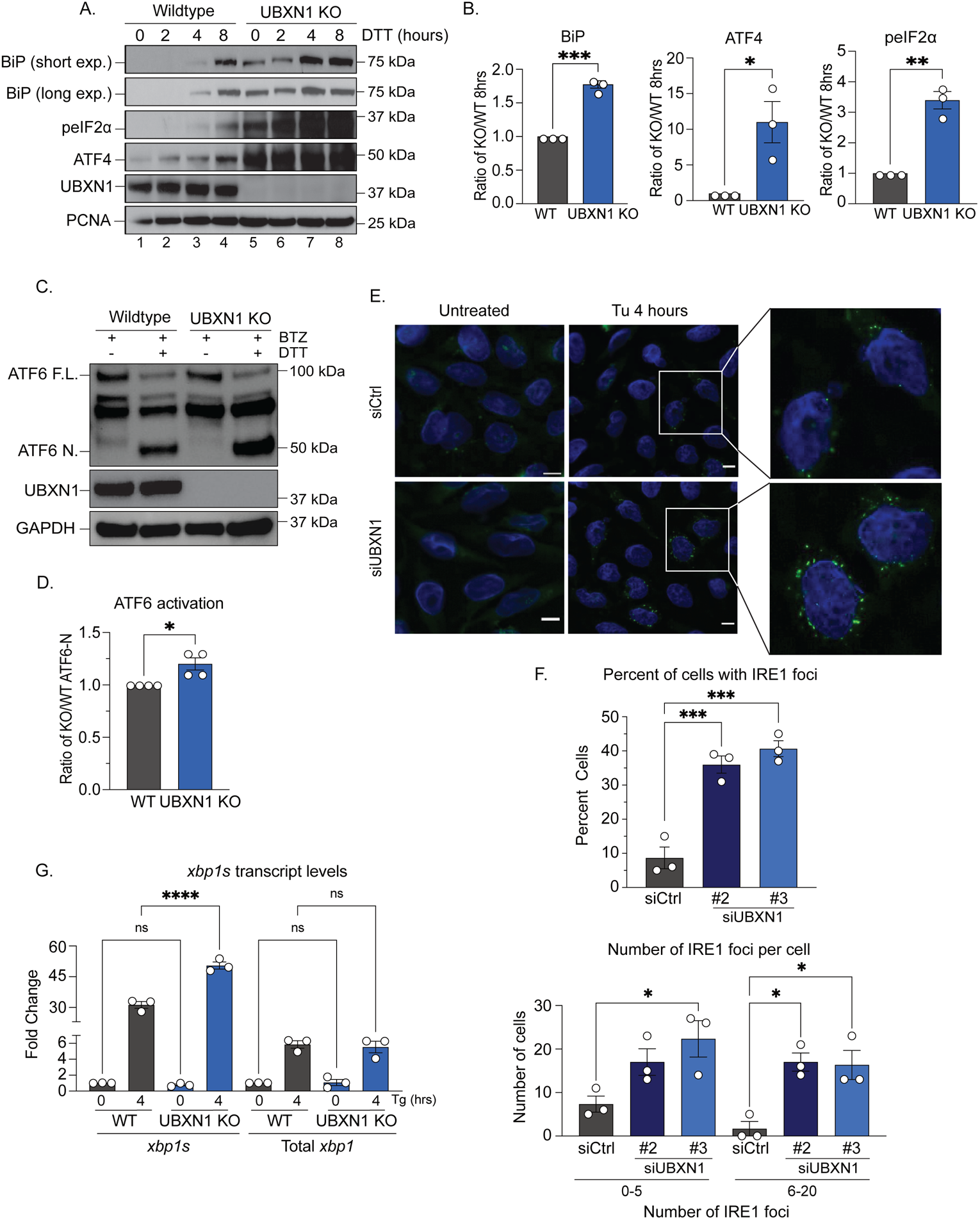
UBXN1 depletion induces activation of the unfolded protein response. **(A)** Immunoblot of HFT wildtype and UBXN1 KO cells treated with 1.5 mM dithiothreitol (DTT) for the indicated timepoints (0-8 hours) to visualize downstream PERK targets. **(B)** Ratio (UBXN1 KO/wildtype) of the band intensity quantifications of BiP, ATF4, and peIF2*α* at the 8-hour timepoint corresponding to Figure 1A. (*n =* three biologically independent samples). We only provide the 8-hour quantification as the signal is not present in wildtype cells in earlier timepoints for accurate quantification. **(C)** Immunoblot of HFT wildtype and UBXN1 KO cells co-treated with 1.5 mM DTT and 1 µM Bortezomib (BTZ) to visualize full-length and cleaved ATF6. **(D)** ATF6 activation was measured by band intensity quantification and calculation of the percentage of cleaved ATF6 to total ATF6. The ratio of the percentage of ATF6 activation in UBXN1 KO cells to wildtype is reported. (*n* = four biologically independent samples). **(E)** Representative immunofluorescent image of IRE1*α* clustering in a stable, doxycycline inducible HFT cell line expressing GFP-tagged IRE1*α*. UBXN1 was depleted by siRNA transfection and cells were treated with 2.5 µM tunicamycin (Tu) for 4 hours to induce IRE1*α* clustering (Scale bar: 10 µm). **(F)** Quantification corresponding to Figure 1E reporting the percent of cells with GFP-IRE1*α*foci as well as the number of GFP-IRE1*α* foci per cell (*n =* three biologically independent samples). **(G)** Transcript levels of *xbp1s* and total *xbp1* in HFT wildtype and UBXN1 KO cells quantified by real-time PCR. Cells were treated with 10 nM thapsigargin (Tg) for 4 hours as indicated. (*n =* three biologically independent samples). Data are means ± SEM (*, **, ***, **** where *P* < 0.05, 0.01, 0.001, and 0.0001, respectively.) Unpaired two-tailed *t* test **(B, D)** or One-way ANOVA with Tukey’s multiple comparisons test **(F, G)**.

Our group recently reported that the p97-UBXN1 complex was required for recognizing ubiquitylated protein aggregates and consolidating them in a perinuclear structure known as the aggresome^54^. ERAD substrates can also be recruited to these aggregates that can associate with the ER membrane to cause ER stress. To determine whether UPR activation caused by UBXN1 loss was due to the accumulation of ubiquitylated aggregates we treated wildtype and UBXN1 KO cells with ER stressors tunicamycin (Tu), which inhibits the first step in N-linked glycosylation, or thapsigargin (Tg), which inhibits the endoplasmic reticulum Ca^2+^ ATPase. Cells were also treated with Bortezomib to induce aggresomes as a positive control. We visualized ubiquitylated proteins by microscopy and immunoblotting. Bortezomib-treated cells contained a single ubiquitin-positive aggresome that was disrupted in UBXN1 KO cells as we previously reported^54^. However, wildtype and UBXN1 KO cells treated with Tu or Tg had no ubiquitin positive aggregates or overt accumulation of ubiquitylated proteins (Supplementary Figure 1G and H). Thus, UPR activation upon UBXN1 deletion is not due to mishandling of aggregates destined for the aggresome. Taken together, our findings implicate UBXN1 as a novel regulator of the UPR independent of ERAD.

### Loss of UBXN1 activates an ER stress-related transcriptional program leading to cell death

The UPR ultimately terminates in a transcriptional response downstream of ATF4, ATF6-N, and xbp1s, therefore, we analyzed UPR target gene expression using quantitative real-time PCR of 84 UPR-specific target genes (Supplementary Table 1). Gene expression analysis in wildtype and UBXN1 KO cells identified a global upregulation (88%, 74 of 84 targets) of UPR-specific target genes in UBXN1 KO cells with 40 genes (47%) experiencing a log_2_ fold change *≥* 0.5 (Figure 2). While this appears to be a modest change, previous microarray analysis has demonstrated that only a handful (22 out of 1868 analyzed) of UPR related genes have 2-fold induction in HeLa cells stressed with tunicamycin^55^. Of the genes that demonstrated a log_2_ fold change *≥* 0.5, there are known targets of each of the UPR sensors, including PERK (*ddit3/chop*), ATF6 (calreticulin, *(calr*) and *herpud1*), and IRE1 (*sec62* and *edem3*), among others^12, 56^. Hierarchical clustering analysis demonstrates that the gene expression pattern observed in UBXN1 KO cells more closely resembles wildtype cells stressed with DTT than untreated wildtype cells based on similar log2 fold change values (Figure 2). Additionally, we observed that the transcriptional response in UBXN1 KO cells treated with DTT is comparable to wildtype cells treated with DTT (Figure 2). We attribute this to a ceiling effect that is often seen with UPR target gene expression upon ER stress^57, 58^.

**Figure 2.**
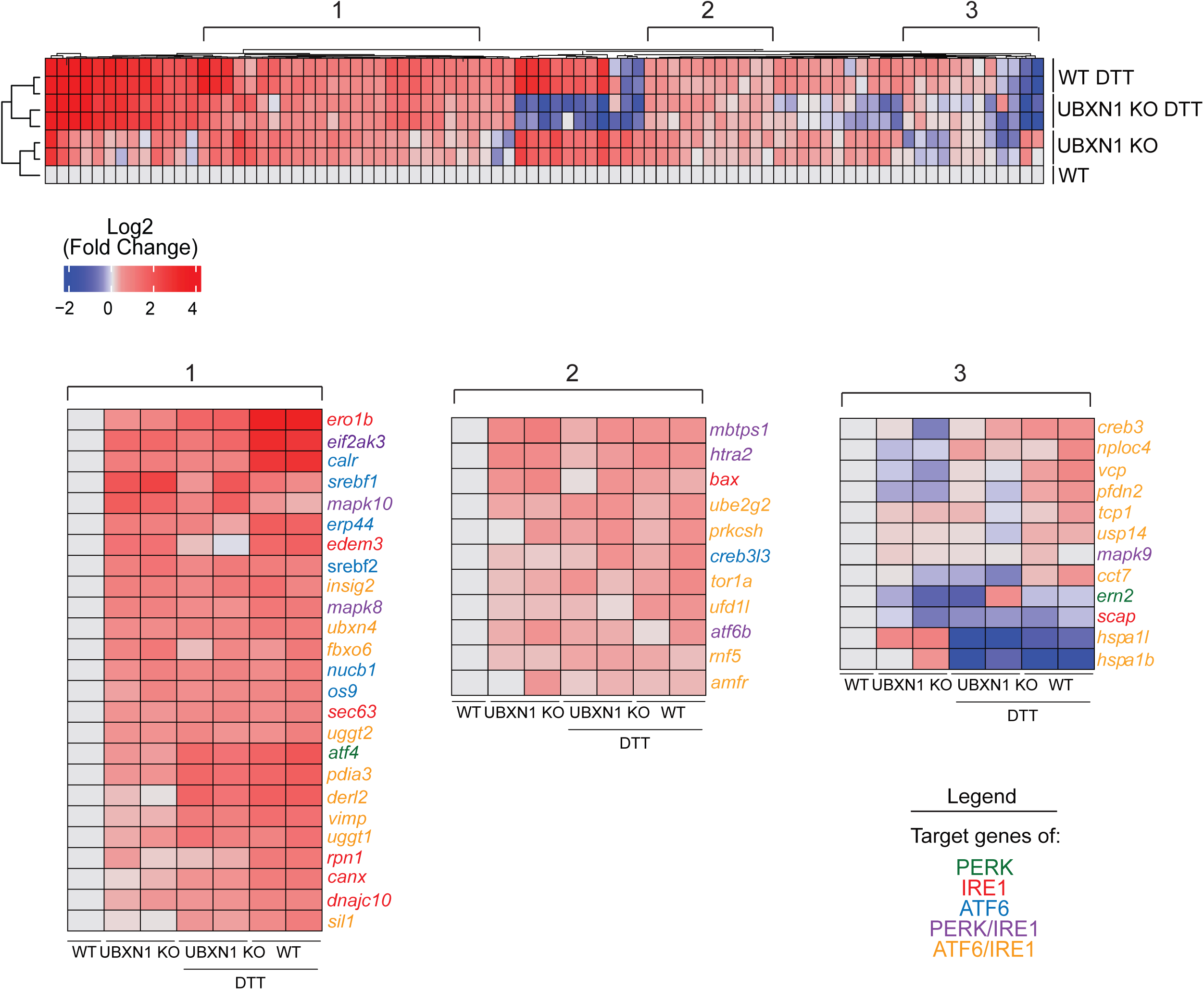
Loss of UBXN1 activates an ER stress transcriptional program. Heat map displays the hierarchical clustering of the log_2_ fold change of 84 UPR target genes determined by the Human Unfolded Protein Response real-time PCR profiler array. The range is from a log_2_ fold change of −2 (blue) to a log_2_ fold change of 4 (red). The untreated wildtype column is displayed as an average of two replicates with the log_2_ fold change set to zero for this column. Additional columns represent two replicates of untreated UBXN1 KO cells, two replicates of UBXN1 KO cells treated with 1.5 mM DTT, and two replicates of HFT wildtype cells treated with 1.5 mM DTT for 4 hours respectively. Section 1 and 2 represent UPR target genes that experience increased expression in the untreated UBXN1 KO cells as well as UBXN1 KO and wildtype cells treated with DTT. Section 3 represents a set of UPR target genes that experience decreased expression in all samples compared to untreated wildtype samples. Genes are color coded by the upstream UPR stress sensor that induces their expression: PERK (green); IRE1 (red); ATF6 (blue); PERK and IRE1 (purple); ATF6 and IRE1 (orange). (*n =* two biologically independent samples).

UPR signaling during acute ER stress is a protective mechanism, whereas chronic ER stress and UPR activation primes cells toward apoptosis^59–62^. We assessed recovery from ER stress in wildtype and UBXN1 KO cells. Cells were treated with thapsigargin for a time-course spanning 0.5 to 3 hours and released into drug-free media (Figure 3A). Viable cells remaining after three days were stained with crystal violet and quantified. We observed significantly reduced growth of UBXN1 KO cells relative to wildtype cells suggesting that elevated UPR signaling primes UBXN1 KO cells for apoptosis (Figure 3A and B). In agreement with this finding, we observed that UBXN1 KO cells have elevated levels of the apoptosis inducing transcription factor C/EBP homologous protein (CHOP) compared to wildtype cells in response to DTT (Figure 3C and D). Thus, UBXN1 loss is detrimental to cell viability in the face of ER stress.

**Figure 3.**
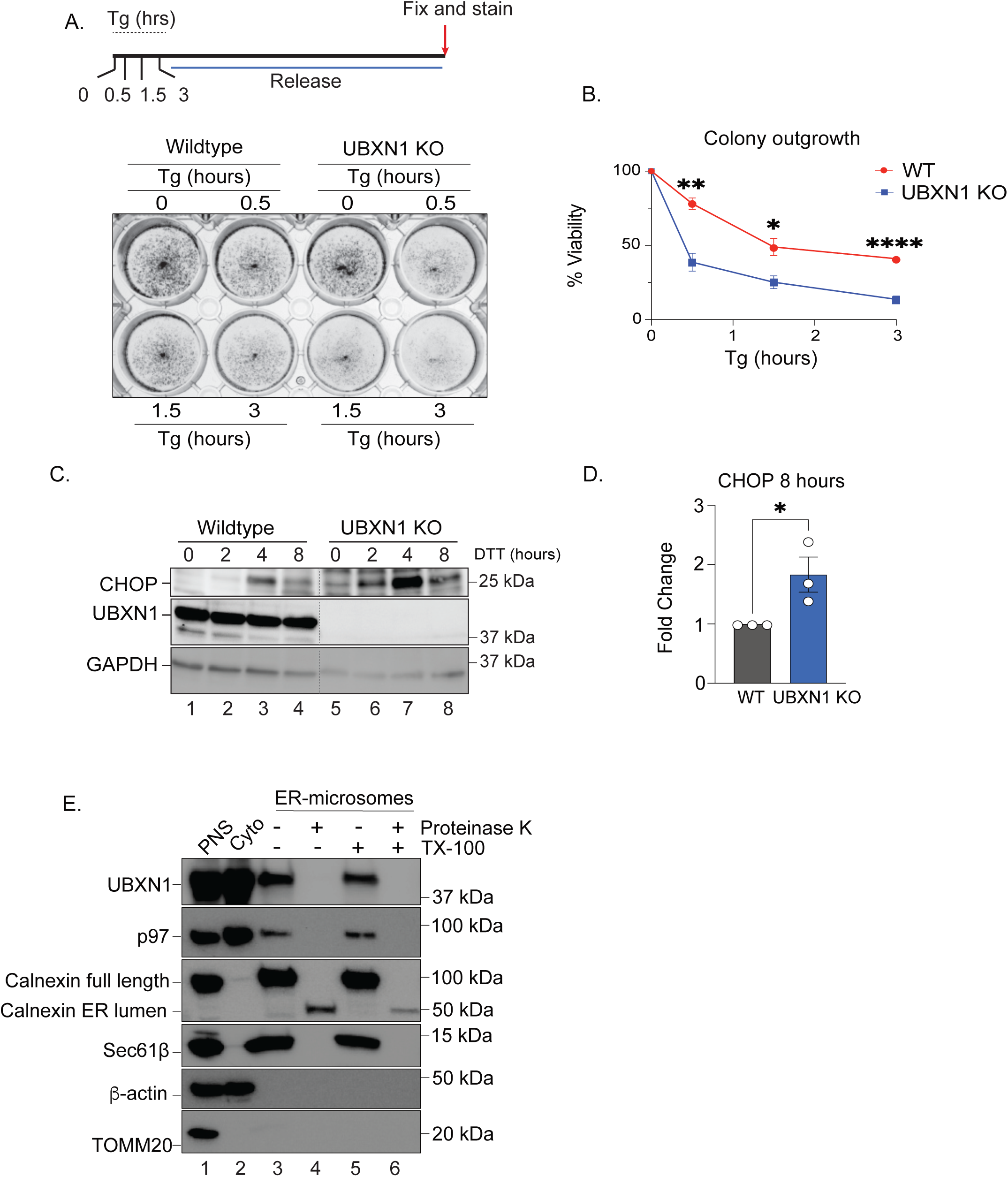
UBXN1 localizes to the ER and preserves cell viability. **(A)** Cell viability of wildtype or UBXN1 KO cells was determined by crystal violet staining. Cells were treated with 200 nM thapsigargin (Tg) for the indicated timepoints and released into drug free media. Colonies were allowed to grow out for three days and live cells were stained with crystal violet. **(B)** Percent viability calculated from the remining cells in Figure 3A. (*n =* four biologically independent samples) **(C)** Immunoblot of HFT wildtype and UBXN1 KO cells treated with 1.5 mM dithiothreitol (DTT) for the indicated timepoints (0-8 hours) to visualize CHOP expression. **(D)** Band intensity quantifications of CHOP expression at the 8-hour timepoint corresponding to Figure 3C. (*n =* three biologically independent samples). We only provide the 8-hour quantification as the signal is not present in wildtype cells in earlier timepoints for accurate quantification. **(E)** Subcellular fractions enriched in ER-derived microsomes were isolated from HEK-293T cells by biochemical fractionation. Protease protection assay was used to localize UBXN1 to the ER periphery by immunoblot. Calnexin and Sec61*β* are ER specific markers, *β*-actin is a cytosolic marker, and TOMM20 is a mitochondrial marker. (PNS: post-nuclear supernatant sample, cyto: cytosolic sample). Data are means ± SEM (*, **, **** where *P* < 0.05, 0.01, and 0.0001, respectively.) Unpaired two-tailed *t* test **(B and D)**

### Loss of UBXN1 leads to a perturbed ER proteome

To understand how UBXN1 loss causes ER stress, we first asked if UBXN1 was associated with the ER. We isolated ER-derived microsomes by biochemical fractionation from HEK293T cells and subjected the microsomes to proteinase K digestion in the presence and absence of detergent (Triton X-100) (Figure 3E). As previously reported, a significant fraction of p97 is peripherally associated with the ER to regulate ERAD^63, 64^ and this localization is sensitive to Proteinase K (Figure 3E). Similarly, we found that UBXN1 was also associated with the ER membrane and was readily digested in Proteinase K treated samples (Figure 3E). We next asked if UBXN1 associated with the SEC61 translocon to regulate import of proteins into the ER, however in co-immunoprecipitation studies we were unable to discern an interaction between UBXN1 and the translocon components SEC61*α* or *β*, whereas UBXN1 interacted with p97 as expected (Supplementary Figure 2A and B).

We decided to take an unbiased approach to assess how UBXN1 deletion impacted ER protein homeostasis. We employed quantitative tandem mass tag (TMT) proteomics. Peptides derived from duplicate wildtype and UBXN1 KO cells were labelled with four-plex TMT labels, combined and peptide abundance was quantified by liquid chromatography and mass spectrometry (Figure 4A). We quantified a total of 6,673 proteins from this study of which 53 were significantly enriched in UBXN1 KO cells (log_2_ fold change KO:WT ≥ 1.0 and p ≤ 0.5) (Figure 4B, Supplementary Table 2). Strikingly, gene ontology analysis identified significant enrichment of ER proteins involved in protein folding, ER-quality control, and the response to ER stress (Figure 4B, C, Supplementary Figure 3A, B). These include proteins involved in *N*-linked glycosylation (glycosyltransferase enzymes ALG1, 3, and 5), ERAD (HRD1 E3 ligase adaptor SEL1L and the ER-anchored protein escortase TMUB1^65^), and ER import (components of the heterotrimeric SEC61 complex: SEC61A1, SEC62, and SEC63 and the ER membrane complex, EMC) (Figure 4C and Supplementary Figure 3A). Notably, when we categorized proteins based on their organelle enrichment, we found that ER proteins in particular were increased in abundance relative to all other organelles (Figure 4D). In contrast, mitochondrial proteins were depleted in UBXN1 KO cells (Figure 4D). Thus, UBXN1 localizes to the ER and loss of UBXN1 alters protein abundance within the ER.

**Figure 4.**
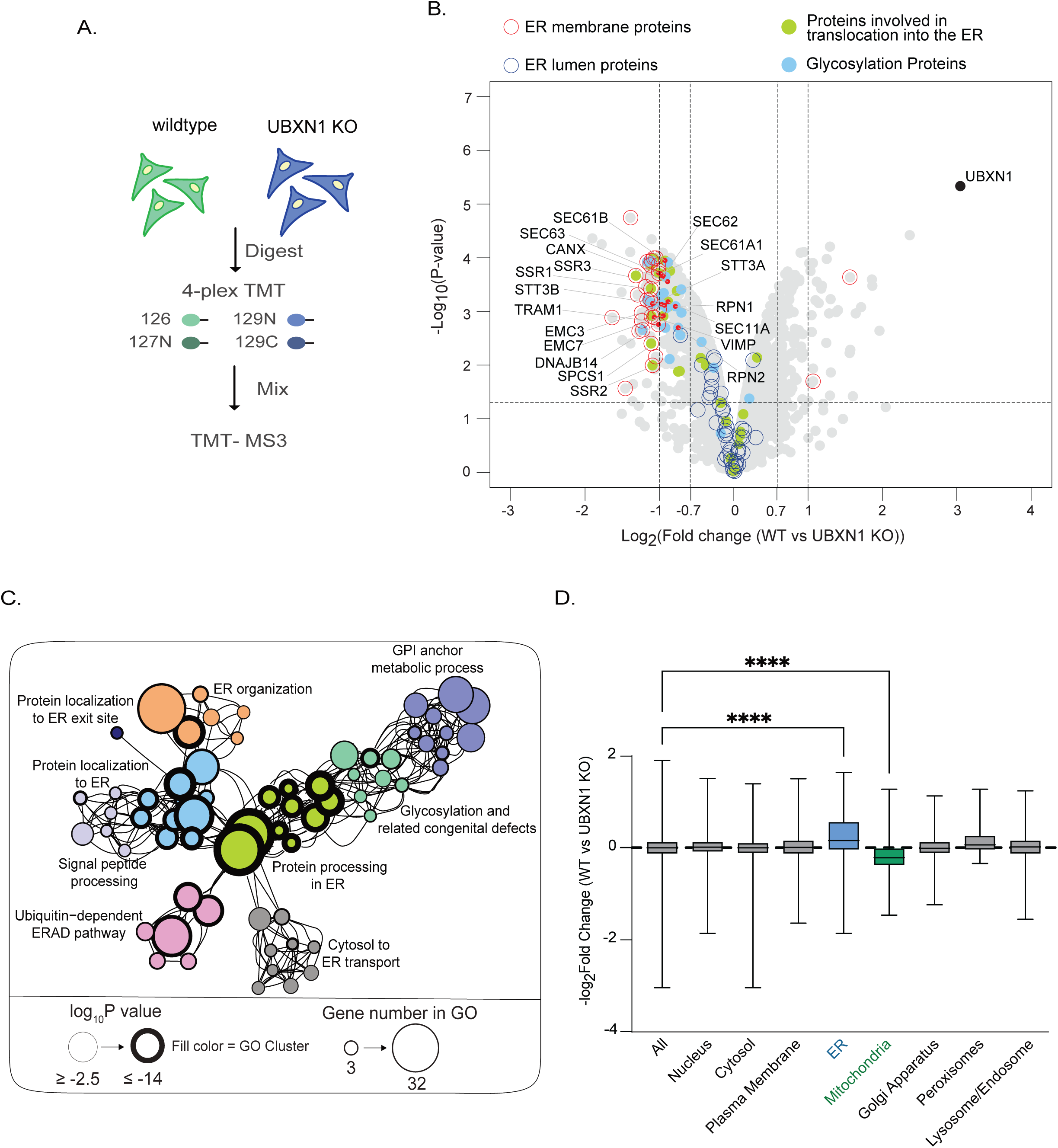
Loss of UBXN1 leads to a perturbed ER proteome. **(A)** Schematic workflow of quantitative tandem mass tag (TMT) proteomics in HFT wildtype and CRISPR-Cas9 UBXN1 knock-out (KO) cells. **(B)** Volcano plot of the (−log_10_-transformed *P* value versus the log_2_-transformed ratio of wildtype/UBXN1 KO) proteins identified by TMT proteomics in wildtype and UBXN1 KO cells. A negative log_2_ fold change indicates proteins with increased abundance in UBXN1 KO cells. Proteins with a log_2_ fold change *≥* |1| are considered significant. ER membrane and luminal proteins are outlined by a red and blue circle respectively. Proteins involved in translocation into the ER and glycosylation are colored in green and blue respectively (*n* = 2 biologically independent samples for each genotype). **(C)** Metascape gene ontology network of the proteins identified to be enriched in UBXN1 KO cells grouped by functional ontology category. The size of the node represents the number of genes in that gene ontology cluster and the thickness of the circle outline corresponds to the −log_10_-transformed *P* value of each term. The different circle colors correspond to a specific gene ontology category. **(D)** Proteins identified in the quantitative proteomics study categorized by organellar compartment. Proteins associated with the ER experience a significant increase in abundance in UBXN1 KO cells where mitochondrial proteins are significantly decreased. Data are means ± SEM (**** where *P* < 0.0001). One-way ANOVA with Brown-Forsythe and Welch ANOVA tests of −logFC **(D)**.

### Depletion of UBXN1 increases the expression and aggregation of ER-destined proteins without preventing their degradation

To further validate our proteomic studies, we examined the expression of significantly enriched proteins by immunoblotting lysates from wildtype and UBXN1 depleted cells. In agreement with the proteomic data, we observed a significant increase in the abundance of the translocon associated protein *α* (TRAP*α*) and alkaline phosphatase placenta-like 2 (ALPP2) (Figure 5A, B, and G). Interestingly, we found that this phenotype extends beyond our proteomics hits; we detected a significant increase in the abundance of ER-targeted proteins *α*-galactosidase (AGAL)^66^ and Prion protein (PrP)^44^ in cells transfected with the corresponding epitope tagged constructs and depleted of UBXN1 (Figure 5C, 5G, Supplementary Figure 4A-C). The elevated levels of these proteins are not due to increased transcription as *prp* transcript levels were unchanged between wildtype and UBXN1 KO cells (Supplementary Figure 4C). We asked if the elevated levels of AGAL and PrP represented ER imported forms or was due to inefficient cytosolic degradation in the absence of UBXN1. We first monitored the glycosylation status of FLAG-AGAL and HA-PrP. AGAL glycosylation was verified by EndoH treatment of cell lysates that resulted in a faster migrating band (Supplementary Figure 4D). HA-PrP glycosylation could be discerned by three distinctly migrating bands representing the unmodified, ER-resident, and fully glycosylated post-ER forms (Supplementary Figure 4A). While UBXN1 depletion resulted in increased steady state levels of these proteins, it did not impact the glycosylation of these proteins indicating that they were targeted appropriately to the ER (Figure 5C, Supplementary Figure 4A, B).

**Figure 5.**
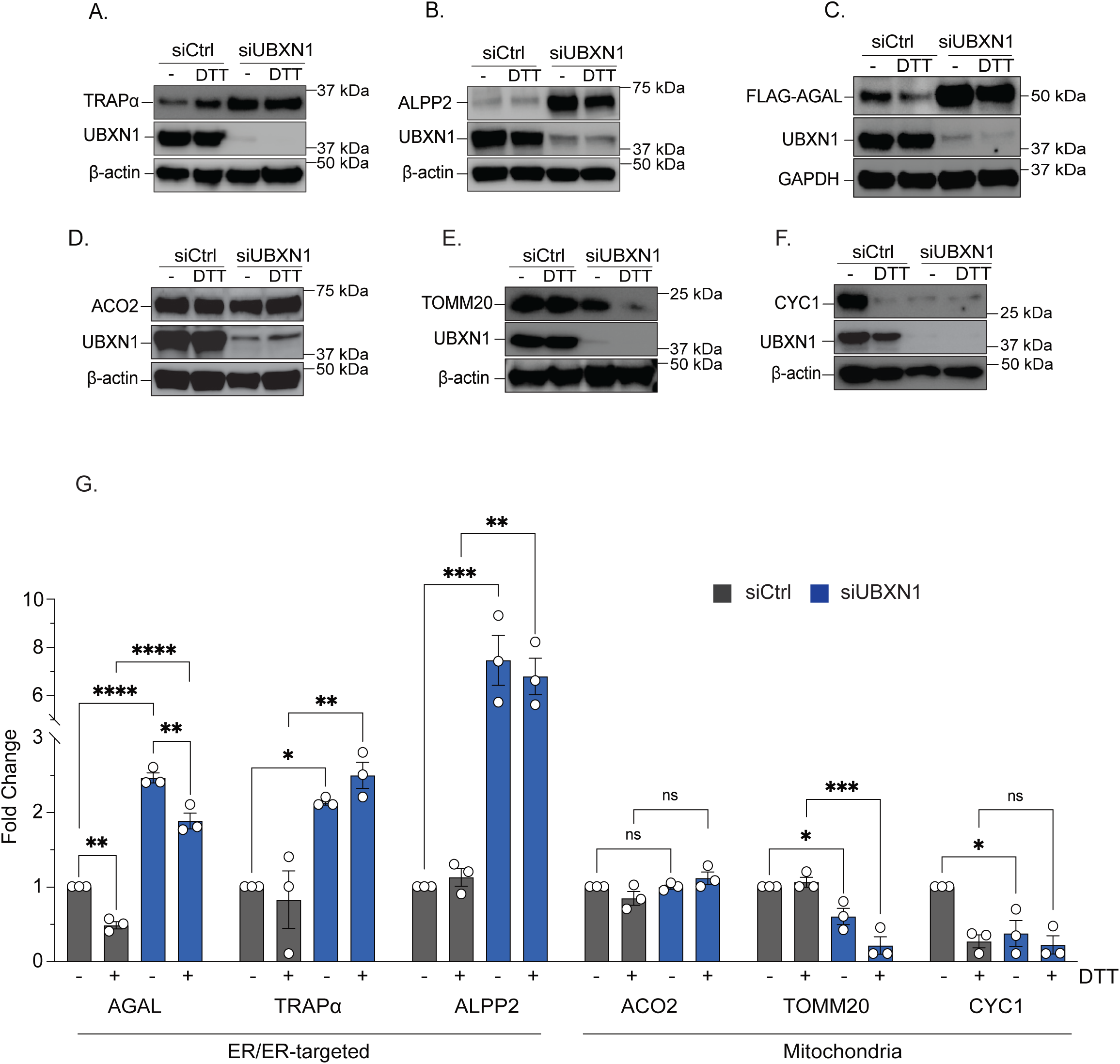
Depletion of UBXN1 increases the expression of ER-destined proteins. **(A and B)** Representative immunoblots of proteins identified to be enriched in the proteomics study. The ER and secretory proteins TRAP*α* and ALPP2 were measured in control cells or cells depleted of UBXN1 for 48 hours with siRNA. Cells were either untreated or treated with 10 mM DTT for 4 hours. **(C)** Immunoblot of a stable HFT cell line expressing doxycycline inducible FLAG-tagged *α*-galactosidase (AGAL). UBXN1 was depleted with siRNA for 24 hours before AGAL induction by 2 µg/ml doxycycline for 24 hours. Cells were untreated or treated with 10 mM DTT for 4 hours. **(D-F)** Representative immunoblots of the mitochondrial resident proteins Aconitase-2 (ACO2), TOMM20, and CYC1 measured in control or UBXN1 depleted cells (48 hours) either untreated or treated with 10 mM DTT for 4 hours. **(G)** Band intensity quantifications of protein expression corresponding to Figure 5A-F. (*n =* three biologically independent samples). Data are means ± SEM (*, **, ***, **** where *P* < 0.05, 0.01, 0.001, and 0.0001, respectively.) One-way ANOVA with Tukey’s multiple comparisons test **(G)**.

Next, we monitored the degradation of HA-PrP and FLAG-AGAL in wildtype and UBXN1 depleted cells. HA-PrP has a weak signal sequence and is mislocalized and degraded prior to import in response to ER stress^44, 46, 67^. This can be visualized by the accumulation of a non-glycosylated cytosolic from in immunoblots (Supplementary Fig 4A). Cells expressing HA-PrP were treated with thapsigargin and the proteasome inhibitor Bortezomib to induce mislocalization and prevent degradation respectively^67^. UBXN1 depletion did not induce greater mislocalization and degradation of HA-PrP relative to total protein (Supplementary Figure 4A compare lane 8 to 4). Furthermore, UBXN1 depletion did not prevent the turnover of FLAG-AGAL or HA-PrP monitored by cycloheximide chase studies (Supplementary Figure 4E-G). UBXN1 depleted cells had significantly higher levels of AGAL compared to control cells to begin with (Figure 5C, 5G, Supplementary Figure 4E), but AGAL was efficiently degraded in both control and UBXN1 depleted cells (Supplementary Figure 4E and F). We note that at earlier timepoints, there was a delay in AGAL degradation in UBXN1-depleted cells and we attribute this to the overall increase in protein abundance of AGAL when UBXN1 is depleted (Figure 5C, 5G). We also assessed the turnover of TRAP*α* and ALPP2 but these proteins had long half-lives and did not appreciably turn over after 9 hours of cycloheximide treatment. Nevertheless, increased protein abundance was obvious (Supplementary Figure 4H).

We asked if increased levels of these protein in the ER impacted their solubility. We monitored AGAL solubility in control and UBXN1-depleted cells treated with DTT and fractionated into soluble and insoluble fractions. In control cells, AGAL was soluble and was not found in the insoluble fraction even upon DTT treatment (Supplementary Figure 4I compare lanes 1 to 3 to 7). Strikingly, in cells depleted of UBXN1 and treated with DTT, AGAL shifted completely into the insoluble fraction (Supplementary Figure 4I compare lanes 2 to 4 and 8).

We were next interested if increased protein abundance upon UBXN1 depletion was unique to ER proteins as suggested by the proteomics (Figure 4D). Our proteomics study identified several mitochondrial proteins depleted in UBXN1 KO cells, but no change in other organellar proteins (Figure 4D). Therefore, we investigated by immunoblot if loss of UBXN1 impacted cytosolic, nuclear, or mitochondrial proteins. While we observed no change in the abundance of proteins in the cytosol or nucleus (Supplementary Figure 4J-M), we observed a significant decrease in the expression of several mitochondrial proteins when UBXN1 was depleted (for example, the mitochondrial outer membrane translocon component TOMM20 and the complex III subunit CYC1) (Figure 5D-G, Figure 4D). Previous studies have demonstrated that the physical interaction between the ER and mitochondria at contact sites facilitates the transmission of ER stress to the mitochondria^68, 69^. To prevent mitochondrial dysfunction transmitted by ER stress, translational attenuation through eIF2α phosphorylation mediates the protective degradation of the TIMM17A subunit of the TIMM23 import complex to prevent protein import into mitochondria^70^. Whether other mitochondrial proteins are similarly downregulated is unknown, but our studies suggest that loss of UBXN1 differentially regulates mitochondrial protein levels in response to ER stress.

Taken together, loss of UBXN1 increases the abundance of ER proteins and leads to their aggregation upon ER stress.

### UBXN1 represses protein translation

Our findings thus far suggested that loss of UBXN1 increased ER protein abundance in a degradation independent manner. We therefore asked if UBXN1 directly impacted protein synthesis. We utilized puromycin incorporation into newly synthesized proteins as a proxy for translation^71^. Puromycin is structurally similar to aminoacyl-tRNAs and can occupy the acceptor (A) site on a ribosome to be incorporated into the nascent polypeptide^72, 73^. Puromycilated proteins are pre-maturely released from the ribosome and can be visualized by immunoblotting with a puromycin antibody^74^. Wildtype and UBXN1 KO cells were briefly labelled (∼30 min) with puromycin, and the levels of puromycilated proteins were quantified. We observed a significant increase in puromycilated proteins in UBXN1 KO cells suggestive of increased translation (Figure 6A, B and Supplementary Figure 5A). We additionally treated cells with DTT or thapsigargin to induce ER stress and shut down protein translation through eIF2α phosphorylation. DTT, thapsigargin, or cycloheximide treatment resulted in a complete shutdown of protein synthesis in wildtype cells as evidenced by the loss in puromycin signal (Figure 6A, compare lanes 7 and 5 to 3, Supplementary Figure 5B compare lanes 3 to 1). However, although significantly reduced in UBXN1 KO cells, translation could still be observed (Figure 6A, B, Supplementary Figure 5A, B). To assess the role of p97 in this process, we performed these studies in cells depleted of p97. Surprisingly, we found that loss of p97 did not impact translation (Figure 6C). In fact, in p97-depleted and thapsigargin treated cells, there was an even more robust termination of translation compared to control cells (Figure 6C and D).

**Figure 6.**
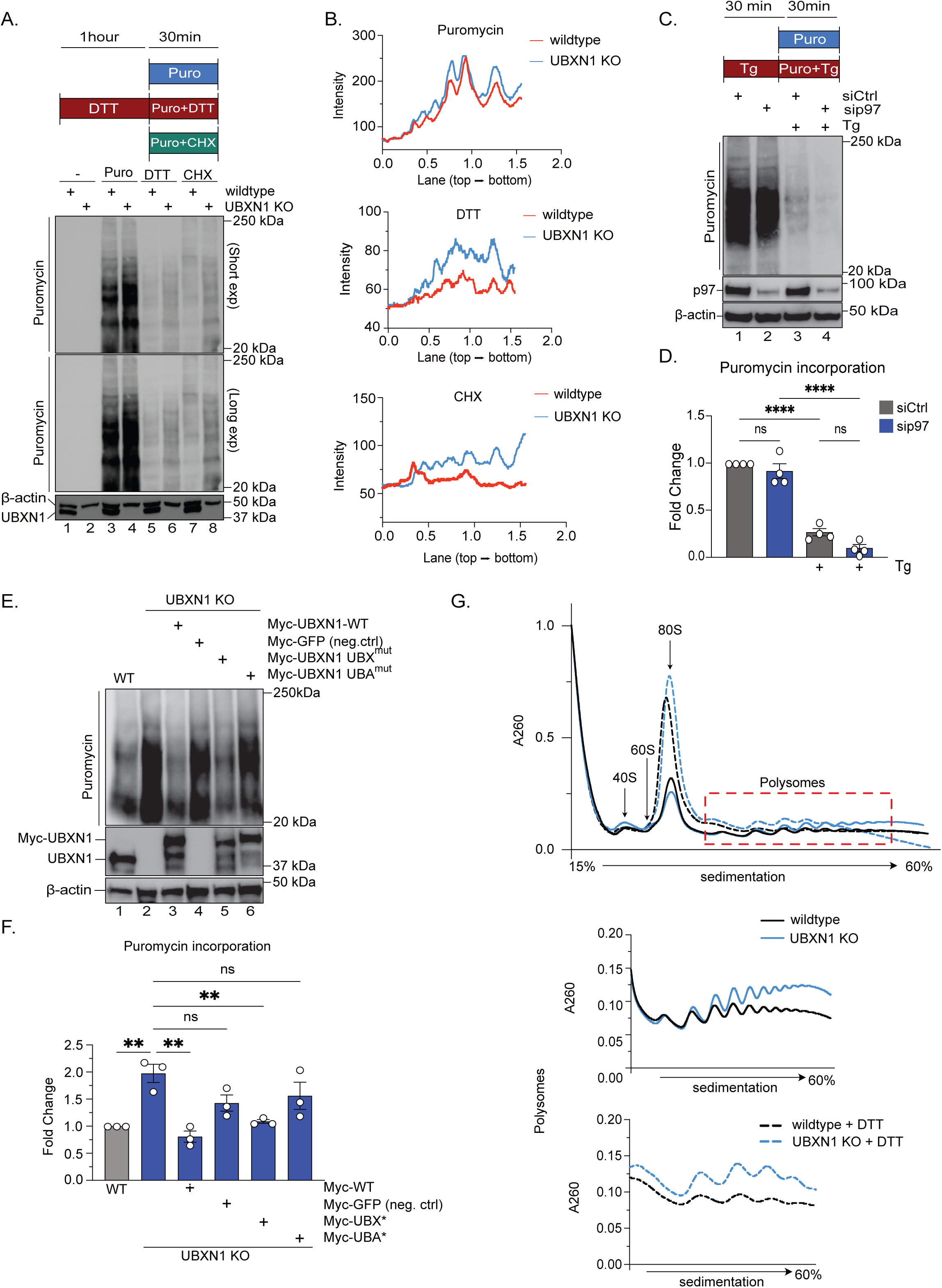
UBXN1 represses protein translation. **(A)** Representative immunoblot of puromycin incorporation into HFT wildtype and UBXN1 KO cells. Cells were pulsed with 1 µM puromycin or 1 µM puromycin in combination with 10 µg/mL cycloheximide (CHX) for 30 minutes. When applicable, cells were pretreated with 1.5 mM DTT for 1 hour before 1 µM puromycin pulse for 30 minutes. The level of puromycin incorporation reflects the rate of protein synthesis. **(B)** Puromycin, DTT, and CHX plots correspond to the intensity of each lane in the immunoblot in Figure 6A. The intensity of the traces for each wildtype and UBXN1 KO sample was determined by the plot profile feature in Fiji. The x-axis represents the distance along the lane. **(C)** Representative immunoblot of 1 µM puromycin incorporation into control cells or cells depleted of p97 with siRNA for 48 hours. Cells were either untreated or pre-treated with 1 µM thapsigargin for 30 minutes. **(D)** Quantification of the whole lane corresponding to each lane in the immunoblot in Figure 6C (*n* = four biologically independent samples). **(E)** Myc-tagged GFP, wildtype, UBX domain mutant, or UBA domain mutant UBXN1 was expressed in cells for 48 hours before 1 µM puromycin pulse for 30 minutes. **(F)** Quantification of each lane intensity from Figure 6E. (*n* = three biologically independent samples) **(G)** Polysome profile traces of HEK-293T wildtype and UBXN1 KO cells either untreated or treated with 2 mM DTT for 60 minutes. The 40S, 60S, and 80S ribosomal subunits are labeled as well as actively translating polysomes. Traces below correspond to the polysomes seen in the main trace. Data are means ± SEM (**** where *P* < 0.0001.) One-way ANOVA with Tukey’s multiple comparisons test **(D, F).**

To determine the specificity of the phenotype, we next asked if we could rescue the perturbed translation phenotype by re-expressing wildtype Myc-UBXN1 in UBXN1 KO cells. As seen in Figure 6E, transfection of Myc-UBXN1 (but not Myc-GFP control) restored translation to wildtype levels (Figure 6E, compare lanes 3 and 4 to lane 1). However, expressing UBXN1 containing mutations in the UBX domain (Phe^265^ Pro^266^ Arg^267^ truncated to Ala-Gly) that we have shown previously to disrupt interaction with p97^54^, also rescued the increase in protein synthesis (Figure 6E and F, compare lanes 5 to 1). However, a UBXN1 UBA domain mutant (Met^13^ and Phe^15^ to Ala) that fails to interact with ubiquitin^40, 54^, did not rescue the increase in protein synthesis to levels observed in wildtype levels (Figure 6E and F compare lanes 6 to 1).

To further confirm the increased translation phenotype, we isolated polysomes to directly analyze the amount of actively translating ribosomes present on mRNA. Inhibiting protein synthesis with cycloheximide halts ribosomes on mRNA such that they can be isolated by sucrose gradient centrifugation and quantified. We found that UBXN1 KO cells have an increase in the levels of actively translating polysomes compared to wildtype cells (Figure 6G). Strikingly, while DTT treatment in wildtype cells halted protein synthesis and collapses polysomes to 80S monosomes, translation persists in UBXN1 KO cells (Figure 6G). In agreement with our puromycin incorporation data, depletion of p97 decreased polysomes in untreated cells that was further depleted upon DTT treatment (Supplementary Figure 5C and D). Taken together UBXN1 represses translation in a p97-independent but ubiquitin-dependent manner (Figure 7).

**Figure 7.**
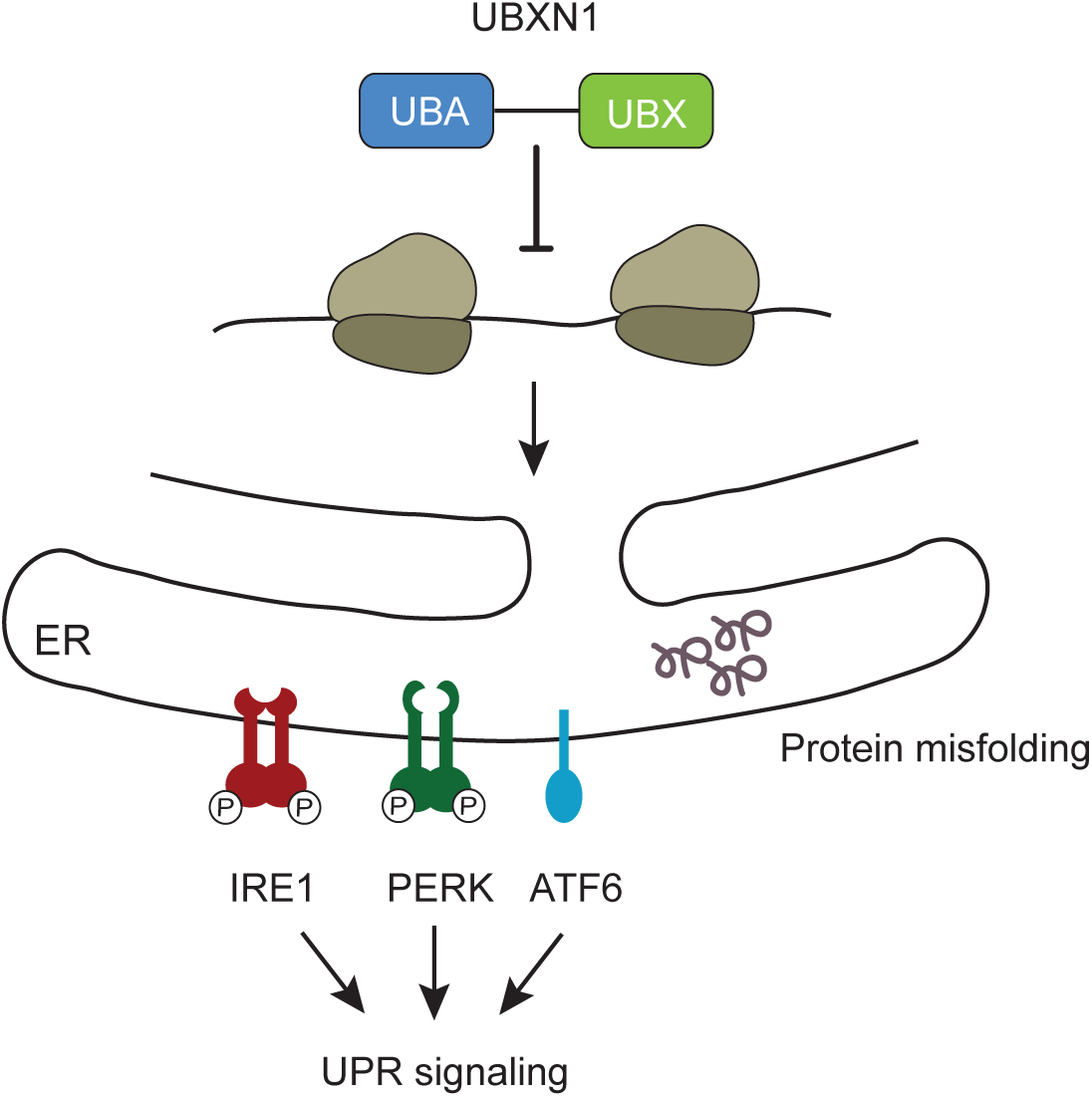
UBXN1 represses protein translation to prevent ER stress and UPR activation. UBXN1 plays an important role in ER-quality control to maintain ER proteostasis. Loss of UBXN1 increases protein synthesis. Increased abundance of ER proteins leads to protein misfolding and activation of the UPR.

## Discussion

Cellular stress can provoke an imbalance between the protein folding machinery and ER-protein load. Numerous studies have illuminated how protein misfolding and consequent ER stress is a major disease mechanism that exacerbates the pathogenesis of a variety of disease states. Failure to adapt to ER stress has been linked to inflammation and UPR-induced apoptosis in cardiovascular disease, liver fibrosis, and diabetes mellitus^75–78^. Thus, it is critical to understand the protein quality control mechanisms that function to prevent ER stress and maintain ER proteostasis to prevent disease.

In the current study, we have identified a novel role for the p97 adaptor protein UBXN1 in ER-quality control as a repressor of the UPR (Figure 7). We found that cells devoid of UBXN1 have significant activation of pathways downstream of PERK, IRE1, and ATF6 (Figure 1) resulting in a transcriptional program to reset ER proteostasis (Figure 2). Our proteomic studies indicate that UBXN1 KO cells have a perturbed ER-proteome with an enrichment of proteins involved in protein translocation into the ER, protein folding, ER-quality control, and the response to ER stress (Figure 4). The increased abundance of these ER proteins is not due to defective degradation via ERAD (Supplementary Figure 4). Furthermore, increased protein abundance in the ER triggers aggregation as observed with AGAL. We propose that the increased influx of proteins into the ER in UBXN1 KO cells causes UPR activation.

A recent study reported that re-initiation of protein synthesis before the restoration of ER proteostasis can induce cell death in response to ER stress^60^. This study found that forced CHOP and ATF4 expression after ER stress induces the expression of several aminoacyl-tRNA synthetases and ribosomal proteins leading to increased translation. Increased protein synthesis elevated reactive oxygen species (ROS) to trigger ER-stress induced apoptosis^60, 61^. Therefore, the ability to attenuate protein synthesis is paramount to cell viability. Indeed, MEFs harboring an S51A mutation in eIF2α which prevents phosphorylation of eIF2α by PERK were significantly sensitized to ER stress, whereas ATF4 null cells, which more slowly restored protein synthesis, were resistant to ER-stress induced cell death^60^. We find robust elevation in the levels of ATF4 (Figure 1) and CHOP (Figure 3) in UBXN1 null cells and these cells are more sensitive to ER stress than their wildtype counterparts (Figure 3).

Finally, we show that UBXN1 is a translation repressor and loss of UBXN1 increases protein translation under basal conditions and in response to ER stress. Whether this occurs through increased ATF4 expression is under investigation. Our study does not address whether ER proteins are specifically repressed by UBXN1 or if it represses translation globally. However, our analysis of the proteomic results and validation studies suggest that ER proteins in particular are more abundant in UBXN1 deleted cells (Figure 4 and 5). How UBXN1 represses translation is currently under investigation.

Surprisingly, we observed that the role of UBXN1 in repressing protein translation is independent of p97 function but reliant on its ability to bind ubiquitin (Figure 6). p97 adaptors are often thought to function exclusively in complex with p97, however, some studies suggest that UBXN1 may function in the absence of p97. For example, after infection by RNA viruses, RIG-I-like receptors in the cytoplasm bind to mitochondrial-antiviral signaling protein (MAVS) to rapidly induce the expression of type I interferons (IFNs)^79^. UBXN1 is recruited to MAVS where it sterically inhibits the binding of downstream effector molecules that contribute to IFNs production^80^. This function was found to be p97 independent as p97 depletion with siRNA, or mutation of the UBX domain did not impact suppression of these pathways. Only the UBA domain was shown to be necessary and sufficient to inhibit antiviral signaling^80, 81^. Whether other adaptors have functions independent of the p97 ATPase remains to be elucidated.

Emerging studies indicate the presence of “pre-emptive” ER-quality control systems that preclude protein misfolding in the ER by preventing the insertion of aberrant proteins. These pathways act upstream of UPR and ERAD. The BAG6-RNF126 mediated degradation of mislocalized ER proteins is one such pre-emptive quality control mechanism^40, 44, 46, 67^. Recently, Ufmylation of the ribosomal protein uL24 (Rpl26) has been reported to enable the degradation of stalled polypeptides in the translocon^82, 83^. It is likely that multiple measures exist to maintain an optimal ER proteome prior to UPR and ERAD activation but they remain to be fully elucidated. We propose that UBXN1-mediated translation repression is one such measure that warrants further investigation.

**Supplementary Figure 1.**
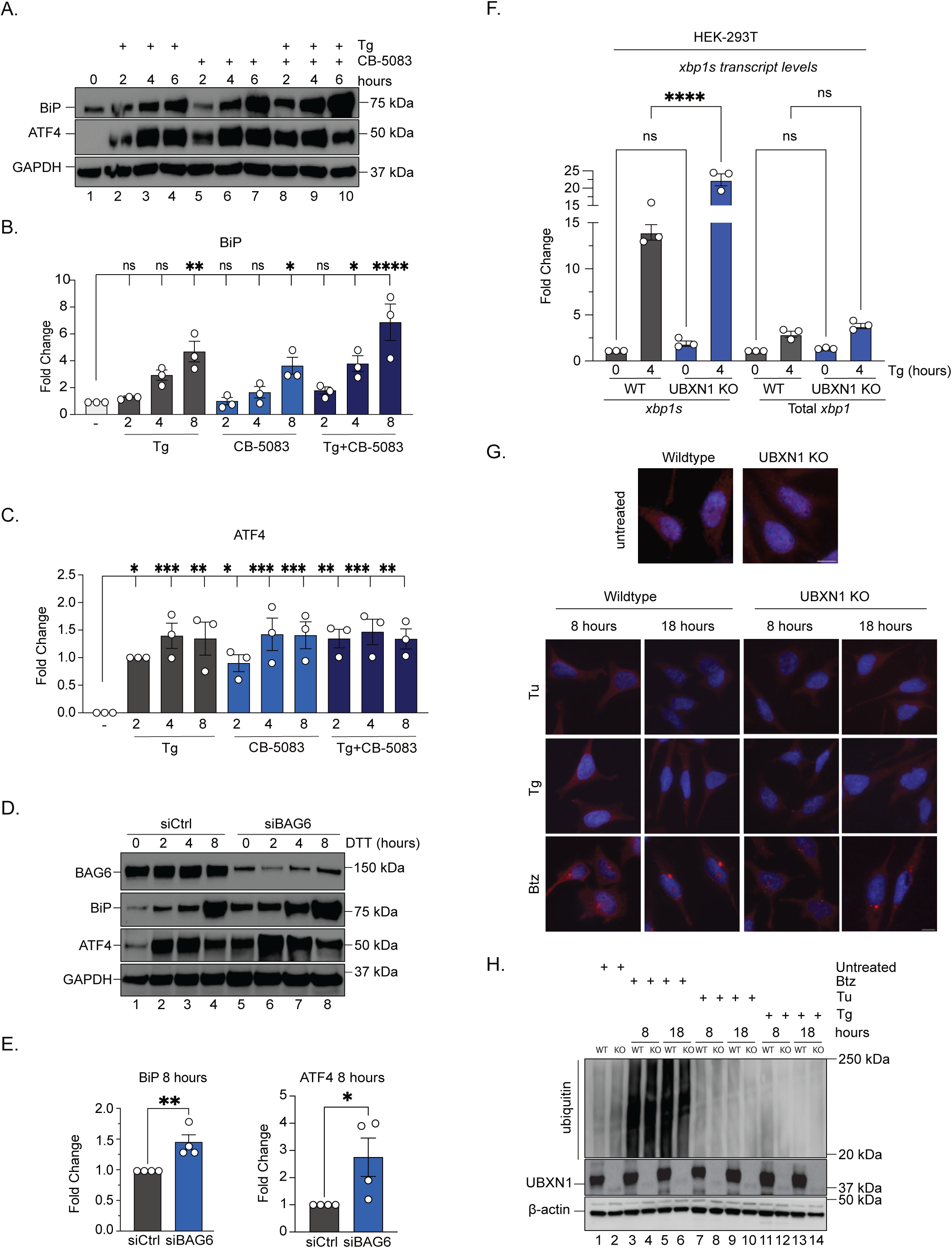
BAG6 depletion and pharmacological inhibition of p97 induces activation of the unfolded protein response. **(A)** Immunoblot of BiP and ATF4 expression levels in HFT wildtype cells treated with 1 µM thapsigargin (Tg), 5 µM of the ATP-competitive p97 inhibitor CB-5083, or both for the indicated timepoints (0-6 hours). **(B-C)** Band intensity quantifications of BiP **(B)** and ATF4 **(C)** corresponding to Supplementary Figure 1A reporting the fold change compared to the untreated condition (*n* = three biologically independent samples)**. (D)** Immunoblot of HFT wildtype cells depleted of BAG6 with siRNA for 48 hours before treatment with 1.5 mM DTT for the indicated timepoints (0-8 hours). **(E)** Band intensity quantifications of BiP and ATF4 corresponding to Supplementary Figure 1D. at the 8-hour timepoint. (*n =* four biologically independent experiments). **(F)** Transcript levels of *xbp1s* and total *xbp1* in HEK-293T wildtype and UBXN1 KO cells quantified by real-time PCR. Cells were treated with 10 nM thapsigargin (Tg) for 4 hours as indicated (*n* = three biologically independent samples). **(G)** Immunofluorescent staining of ubiquitin (FK2) and nuclei (Hoechst dye) in HFT wildtype and UBXN1 KO cells. Cells were treated with 1 µM of the proteasome inhibitor bortezomib (BTZ), Tunicamycin (Tu), or Thapsigargin (Tg) for 8 hours or 1 µM BTZ, 500 nM Tu, or 500 nM Tg for 18 hours. **(H)** Immunoblot assessing total ubiquitin levels corresponding to Supplementary Figure 1G. Data are means ± SEM (*, **, ***, **** where *P* < 0.05, 0.01, 0.001, and 0.0001, respectively.) One-way ANOVA with Dunnetts’s multiple comparisons test **(B and C).** Unpaired two-tailed *t* test **(E).** One-way ANOVA with Tukey’s multiple comparisons test **(F)**.

**Supplementary Figure 2.**
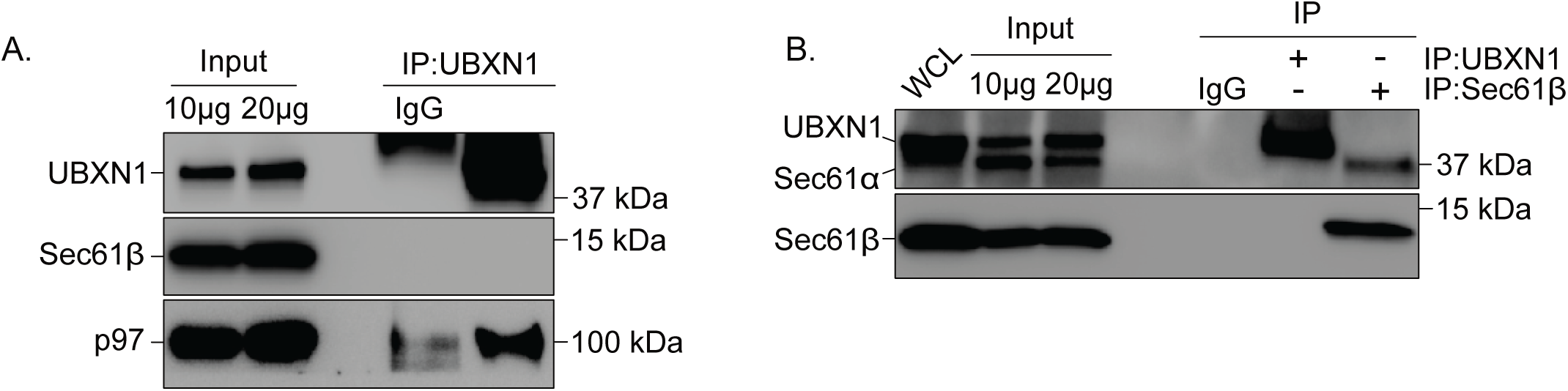
UBXN1 does not interact with the Sec61 translocon. **(A)** Representative immunoblot for the immunoprecipitation of endogenous UBXN1 from HEK-293T lysates. 10 and 20 µg of whole cell lysate was used for the input. Sec61*β* was used as a marker for the Sec61 translocon. **(B)** UBXN1 and Sec61*β* were separately immunoprecipitated from ER-microsomes isolated from HEK-293T cells. 10 and 20 µg of whole cell lysate was used for the input.

**Supplementary Figure 3.**
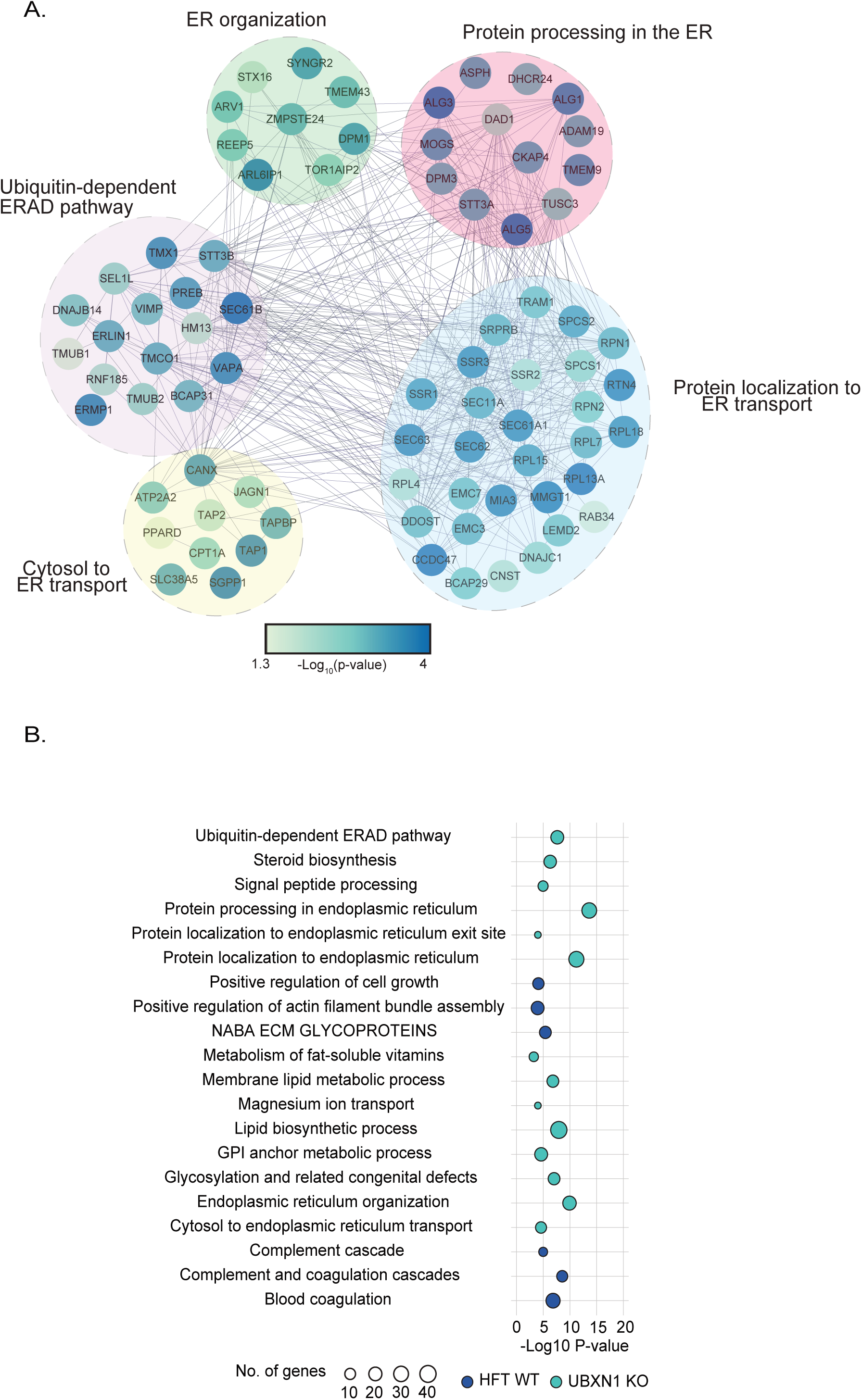
Loss of UBXN1 leads to a perturbed ER proteome. **(A)** Interaction network of the gene ontology clusters from the wildtype/UBXN1 KO quantitative proteomics study with a significant increase in abundance in UBXN1 KO cells. Proteins found in each node are labelled and the color of each circle corresponds to the −log_10_-transformed *P* value. **(B)** Bubble plot corresponding to significantly enriched gene ontology groups in the wildtype (navy) and UBXN1 KO (teal). The size of the circle corresponds to the number of genes identified in each term.

**Supplementary Figure 4.**
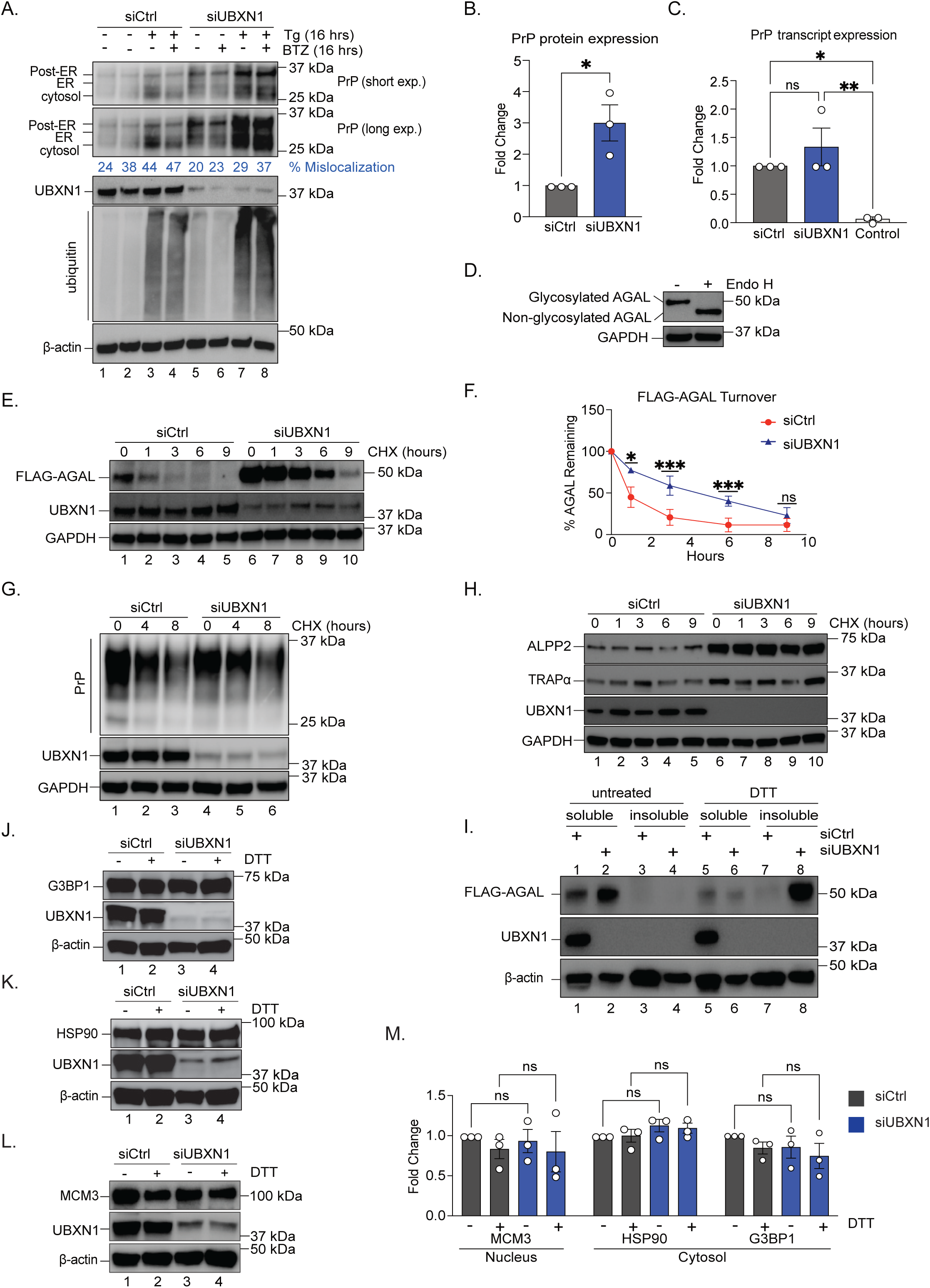
Depletion of UBXN1 increases the expression of ER-destined proteins. **(A)** Immunoblot of HFT wildtype cells expressing HA-tagged prion protein (PrP). PrP was expressed in HeLa cells for before siRNA mediated depletion of UBXN1. Cells were treated with 500 nM thapsigargin, 1 µM bortezomib (BTZ), or both for 16 hours. The percent mislocalization was calculated from the ratio of the mislocalized (cytosolic) form to all forms of PrP for that sample. **(B)** Total PrP protein expression was calculated by the sum of all forms of PrP for that sample. The untreated condition was quantified (lanes 1 and 5) (*n* = three biologically independent experiments). **(C)** Transcript levels of HA-PrP in cells depleted of UBXN1 quantified by real-time PCR. Control is cDNA generated from cells not expressing HA-PrP. (*n =* three biologically independent experiments). **(D)** Immunoblot of a stable HFT cell line expressing doxycycline inducible FLAG-tagged AGAL. Glycans were digested with EndoH for 4 hours to generate the non-glycosylated form. **(E)** Immunoblot examining the turnover of FLAG-AGAL in siRNA control and UBXN1 depleted cells (48 hours). Cells were treated with 10 µg/mL cycloheximide (CHX) for the indicated timepoints (0-9 hours). **(F)** Band intensity quantification of each time point of the CHX time course corresponding to Supplementary Figure 5E. The zero hour time point was set to 100 % for each siRNA and the % of AGAL remaining for each time point was quantified as a ratio from the zero hour timepoint for that siRNA. (*n ≥* three biologically independent samples. **(G)**Immunoblot examining the turnover of HA-PrP in siRNA control and UBXN1 depleted cells. Cells were treated with 10 µg/mL cycloheximide (CHX) for the indicated timepoints (0-8 hours). **(H)**Immunoblot examining the turnover of the TMT-proteomics hits (ALPP2 and TRAP*α*) that were identified to be significantly enriched in cells depleted of UBXN1. Control and UBXN1 depleted cells were treated with 10 µg/mL cycloheximide (CHX) for the indicated timepoints (0-9 hours). **(I)** Immunoblot of soluble and insoluble fractions of FLAG-tagged AGAL in control cells or cells depleted of UBXN1 with siRNA. Cells were either untreated or treated with 10 mM DTT for 4 hours. (In this experiment only, we quantified and ran 2.5x less siUBXN1 protein so we could accurately visualize and compare siControl to siUBXN1. The AGAL expression in siUBXN1 cells often bleaches out what can be observed with siControl.) **(J-L)** Immunoblots of samples from Figure 5A. examining the protein expression of additional organellar proteins including the nuclear protein MCM3 and the cytosolic proteins G3BP1 and HPS90. **(M)** Band intensity quantifications of protein expression corresponding to Supplementary Figure 5J-L. (*n =* three biologically independent samples). Data are means ± SEM (*, **, and *** where *P* < 0.05, 0.01, and 0.001 respectively.) Unpaired two-tailed *t* test **(B and F)**, One-way ANOVA with Tukey’s multiple comparisons test **(C and M)**.

**Supplementary Figure 5.**
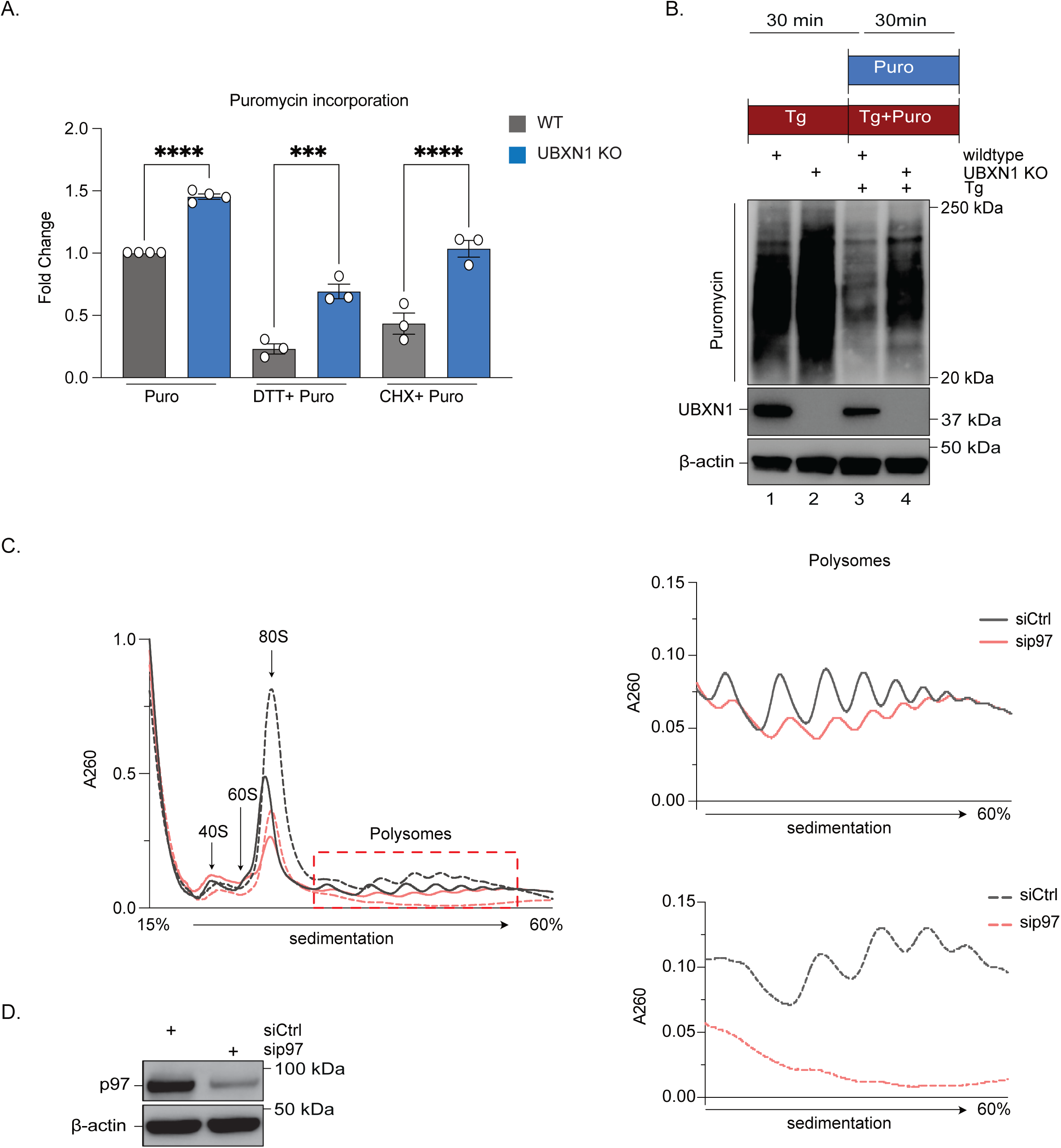
UBXN1 suppresses protein synthesis. **(A)** Densitometry quantifications of the entire lanes corresponding to Figure 6A (*n ≥*three biologically independent experiments). **(B)** Immunoblot of HFT wildtype and UBXN1 KO cells pulsed with 1 µM puromycin for 30 minutes after pre-treatment with 1.5 µM thapsigargin for 30 minutes. **(C)** Polysome profile traces of HEK-293T control cells or cells depleted of p97 with siRNA for 72 hours. Cells were either untreated or treated with 2 mM DTT for 60 minutes. The 40S, 60S, and 80S ribosomal subunits are labeled as well as actively translating polysomes. Traces on the right correspond to the polysomes seen in the traces on the left. **(D)** Immunoblot validating the siRNA knockdown corresponding to Supplementary Figure 6C. Data are means ± SEM (*** and **** where *P* < 0.001 and 0.0001, respectively.) One-way ANOVA with Tukey’s multiple comparisons test **(A)**.

## MATERIALS AND METHODS

### Cell culture, transfections, immunoblot, and immunoprecipitation

HeLa Flp-IN T-REX (HFT) and HEK-293T cells were cultured in Dulbecco’s modified Eagle’s medium (DMEM) supplemented with 10 % fetal bovine serum (FBS) and 100 U/mL penicillin and streptomycin. Cells were grown in a humidified, 5 % CO_2_ atmosphere at 37 °C. For siRNA transfections, cells were either forward or reverse transfected with 20 nM siControl, siUBXN1, siBAG6, or sip97 using Lipofectamine RNAiMax (Invitrogen) in 6-well plates. After 8 or 24 hours depending on the study, cells were split into 12-well plates for treatment. 48 hours post transfection, cells were treated with the indicated drugs for the indicated timepoints before harvest for immunoblot or fixation for immunofluorescence. For DNA transfections, 1 µg HA-prion protein (PrP), 0.5 µg Myc-tagged wildtype UBXN1, 0.5 µg of either Myc-tagged UBX mutant (Phe^265^ Pro^266^ Arg^267^ truncated to Ala-Gly), or Myc-tagged UBA mutant (Met^13^ and Phe^15^ to Ala), and 50 ng Myc-tagged GFP constructs were forward transfected into HFT cells seeded in a 6-well plate using Lipofectamine 3000 (Invitrogen). The cells were harvested 48 or 72 hours post transfection. To establish an HFT cell line stably expressing doxycycline inducible FLAG-AGAL, 1 µg FLAG-AGAL construct and 4.5 µg pOG44 was transfected into HFTs seeded onto a 6-well plate using Lipofectamine 2000 (Invitrogen). After 48 hours the transfected cells were selected with hygromycin (200 µg/mL) for about 10 days.

Cell pellets were lysed in either radioimmunoprecipitation assay (RIPA) buffer (25 mM Tris-Cl [pH 7.6], 150 mM NaCl, 1 % NP-40, 1 % sodium deoxycholate, 0.1 % SDS, HALT protease inhibitors (Pierce PI-78425), sodium vanadate, sodium fluoride) or in 1 % SDS lysis buffer (1 % SDS, 50 mM Tris-Cl, 150 mM NaCl, 0.5 % NP-40, HALT protease inhibitors, sodium vanadate, sodium fluoride). FLAG-AGAL and HA-PrP cell pellets were resuspended in RIPA buffer and incubated on ice for 20 minutes with a 10 second vortex pulse every 5 minutes. Lysates were then centrifuged at 14,000 rpm for 15 minutes at 4 °C. For UPR induction studies in Figure 1, cell pellets were resuspended in SDS lysis buffer, vortexed briefly, boiled for 10 minutes at 95 °C, and briefly vortexed. Lysates were incubated at 65 °C for an additional 5 minutes before centrifugation at 14,000 rpm at room temperature. The supernatant was collected, and protein concentration was estimated utilizing the DCA protein assay kit (Biorad). Protein expression was analyzed by SDS-PAGE and immunoblot.

For immunoprecipitations, cell pellets were lysed in mammalian cell lysis buffer (50 mM Tris-HCl [pH 7.6], 150 mM NaCl, 1 % Nonidet P-40, HALT protease inhibitors (Pierce), and 1 mM DTT). Protein G agarose beads (Thermo Scientific 20398) was pre-washed with MCLB before addition of the cell lysate and antibody (UBXN1, Sec61*β*, or rabbit IgG control). Samples were incubated with rotation overnight at 4 °C. Beads was collected by centrifugation at 3000 rpm for 1 minute and was washed four times with MCLB buffer. Beads were resuspended in 50 µl SDS sample buffer, briefly boiled, and processed for SDS-PAGE and immunoblotting.

### Antibodies, siRNA, and reagents

The rabbit UBXN1 (16135-1-AP 1:7000 dilution), p97 (10736-1-AP), BAG6 (26417-1-AP), ATF6 (24169-1-AP), Sec61*β* (51020-2-AP 1:3000 dilution), Aconitase 2 (11134-1-AP), calnexin (10427-2-AP), CYC1 (10242-1-AP), and G3BP1 (13057-2-AP) antibodies were obtained from Proteintech Inc; rabbit BiP (3177 dilution), peIF2*α* (3398 1:500 dilution), and MCM3 (4012S) antibodies were obtained from Cell Signaling Technology; mouse PCNA (sc-56 1:3000 dilution), ATF4 (sc-390063 1:500 dilution), Tom20 (sc-17764), ubiquitin (P4D1; sc8017), *β*-actin (sc-69879 1:3000 dilution), GAPDH (sc-47724 1:3000 dilution), c-myc (sc-40 1:3000 dilution), Sec61*α* (sc-393182 1:500 dilution), and TRAP*α* (sc-373916) were from Santa Cruz Biotechnology; mouse ubiquitin FK2 (04-263 1:100 dilution) for immunofluorescence and Anti-Puromycin, clone 12D10 (MABE343 1:3000 dilution) was from EMD Millipore; mouse FLAG-M2 antibody (F3165 1:1000 dilution) was from Sigma Aldrich. The rabbit anti-Prion protein PrP antibody (ab52604) was purchased from Abcam. The rabbit ALPP/ALPP2 was purchased from St. John’s Laboratory (STJ96740). The mouse HSP90AB1 was purchased from Origene (TA500494). All antibodies were used at a dilution of 1:1000 unless otherwise specified. Secondary antibodies, HRP conjugated anti-rabbit (W4011) and anti-mouse (W4021) were from Promega and used at a dilution of 1:10000. Thapsigargin (5860051MG), Tunicamycin (ICN15002801), Proteinase K (FEREO0491), and Hoechst (51-17 1:10000 dilution) for immunofluorescence were purchased from Fisher Scientific. Dithiothreitol (DTT25) and puromycin (P-600-100) were purchased from GoldBio. Cyclohexmide (97064-724) and Triton X-100 (97063-864) was purchased from VWR. CB-5083 were purchased from Cayman Chemical Company (19311). Bortezomib (S1013) was purchased from Selleckchem. EndoH (P0702S) was purchased from New England Biolabs. Crystal violet was purchased from Sigma-Aldrich (C6158-50G). Doxycycline was purchased from EMD Millipore (324385-1GM). Wild-type PrP (HA) was generously gifted by Ramanujan Hegde (MRC, UK). FLAG-tagged AGAL were generously gifted by Malaiyalam Mariappan^66^. siRNAs against UBXN1-2 (D-008652-02), UBXN1-3 (D-008652-03), and UBXN1-4 (D-008652-04) were purchased from Dharmacon. siRNA against BAG6 (s15467) was purchased from Ambion (Themo Fisher Scientific). siControl (SIC001) was from Millipore Sigma. siRNA against p97 was purchase from Invitrogen (HSS111263).

### Generation of CRISPR gene knockout cell lines

UBXN1 knockout cells were generated in both HFT and HEK-293T cells using the CRISPR-Cas9 gene editing method. The guide sequence 5′-GCCGTCCCAGGATATGTCCAA-3′ was cloned into the pX459 vector carrying hSpCas9 as previously published^40^. 500 ng of the construct was transiently transfected into HFT and HEK-293T cells seeded in a 6-well plate with Lipofectamine 3000. 36 hours post-transfection, the cells were cultured with 1 µg/mL puromycin for an additional 24 hours. The surviving cells were counted and serially diluted to a concentration of 1 cell per well in a 96-well plate. Protein expression of UBXN1 was examined by immunoblot.

### Crystal violet cell viability assay

Wildtype and UBXN1 KO cells were plated at a density of 8000 cells/well in a 12-well dish. The next day, cells were treated with 200 nM thapsigargin for the indicated timepoints. After treatment, wells were washed twice with PBS before re-addition of fresh media. Every day for three days, cells were washed once with PBS and media was replaced. On the third day, cells were washed twice with PBS before incubation with 0.5 % crystal violet staining solution for 20 minutes at room temperature on an orbital shaker. After staining, wells were washed thrice with PBS to remove unbound stain. 1 % SDS-solution was added to each well to solubilize bound stain and the plate was incubated for one hour at room temperature on an orbital shaker until color was uniform in solution. The absorbance was read at 570 nm on a multiwell plate reader (SpectraMax iD3). The average absorbance of wells without cells was used to background subtract.

### Real-time PCR

For all real-time PCR experiments, total RNA was isolated using the *Quick-*RNA Miniprep Kit (Zymo Research cat. no. R1055). The purified RNA was quantified by NanoDrop and 1 µg of RNA for each sample was used to generate cDNA using the iScript cDNA synthesis kit (Biorad cat. no. 1708890). Real-time PCR was performed with PowerUp SYBR Green Master Mix (Applied Biosystems cat. no. A25741) on an Applied Biosystems StepOnePlus real-time PCR system. Data analyses utilized the 2^-ΔΔC*t*^ method and GAPDH was used as a housekeeping gene to normalize transcript expression across samples. The XBP1s primers were previously published^52^.

XBP1s forward: 5′ TGCTGAGTCCGCAGCAGGTG 3′
XBP1s reverse: 5′ GCTGGCAGGCTCTGGGGAAG 3′
Total XBP1 forward: 5′ AAACAGAGTAGCTCAGACTGC 3′
Total XBP1 reverse: 5′ TCCTTCTGGGTAGACCTCTGGGAG 3′
HA-PrP forward: 5’ CAGCCTCATGGTGGTGGCTG 3’
HA-PrP Reverse: 5’ GTCGGTCTCGGTGAAGTTCTCC 3’

### RT^2^ Profiler PCR array

Wildtype or UBXN1 KO HeLa Flp-in T-REX cells were grown in 10cm tissue culture dishes and either left untreated or treated with 1.5 mM DTT for 4 hours. Post-harvest, RNA was extracted from each sample with the *Quick*-RNA MiniPrep Kit (Zymo Research cat. no. R1055) and cDNA was generated with the RT^2^ First Strand Kit (Qiagen cat no. 330401). Each Human Unfolded Protein Response RT^2^ Profiler PCR Array (cat no. 330231 PAHS-089ZA) plate consists of 84 UPR specific target genes as well as a panel of housekeeping genes and PCR efficiency controls. Each sample was distributed to an individual plate for two biological replicates each. The GeneGlobe Data Analysis Center was used to determine the fold change between samples. The data were normalized by automatic selection from the housekeeping gene panel utilizing the geometric mean. The untreated wildtype HFT samples were defined as the “control” sample for the data set and used to calculate fold changes. Heatmap visualizations for PCR array data were performed using the RStudio software (v2022.07.1 Build 554), including ggplot2 (v 3.3.5), circlize (v0.4.14), ComplexHeatmap version 2.10.0, and InteractiveComplexHeatmap package to export heatmap generated by ComplexHeatmap into an interactive Shiny app^84^.

### Immunofluorescence and microscopy

To assess aggresome formation, cells were grown on coverslips (12mm) in a 24-well plate treated with 1 µM bortezomib (Btz), Tunicamycin (Tu), or Thapsigargin (Tg) for 8 hours or 1 µM Btz, 500 nM Tu, or 500nM Tg for 18 hours. Cells were washed briefly in phosphate-buffered saline (PBS) and fixed with ice-cold methanol for 10 min at −20°C. Cells were washed in PBS then blocked in 2 % bovine serum albumin (BSA) with 0.3 % Triton X-100 in PBS for 1 hour. The coverslips were incubated with the indicated antibodies in a humidified chamber overnight at 4 °C. Coverslips were washed and incubated with the appropriate Alexa Fluor-conjugated secondary antibodies (Molecular Probes) for 1 hour in the dark at room temperature. Cells were washed with PBS, and nuclei were stained with Hoechst 33342 dye and mounted onto slides. All images were collected using a Nikon A1R scan head with a spectral detector and resonant scanners on a Ti-E motorized inverted microscope equipped with a 60× Plan Apo 1.4 NA objective lens. The indicated fluorophores were excited with either a 405-nm, 488-nm or 594-nm laser line. Images were analyzed by using FIJI (https://imagej.net/Fiji).

### IRE1*α*-GFP foci imaging and quantification

Stable HFT wildtype cells expressing doxycycline inducible IRE1*α*-3xFLAG-6xHis-GFP (construct a generous gift from Peter Walter, UCSF) were seeded onto a 6-well plate and reverse transfected with an siControl or siUBXN1^51^. Next day, cells were trypsinized and seeded onto coverslips in a 12-well plate. 48 hours after siRNA transection, IRE1*α*-GFP was induced with 4 µg/mL doxycycline for an additional 24 hours. Next day, cells were either left untreated or treated with 2.5 µM tunicamycin for 4 hours. Coverslips were briefly washed with phosphate-buffered saline (PBS) and fixed with ice-cold methanol for 10 min at −20 °C. Coverslips were washed with PBS and nuclei were stained with Hoechst 33342 dye and mounted on slides. Images were collected using either a 405-nm or 488-nm laser. One hundred cells were counted by creating a cell mask and utilizing the “Find Maxima” function in FIJI (https://imagej.net/Fiji).

### Subcellular fractionation of ER-derived microsomes

HEK-293T cells were seeded into four 150 mm sterile tissue culture dishes. Cells were scraped off the plate into PBS, centrifuged at 600 x *g* for 5 minutes at 4 °C, and resuspended in ice-cold homogenization buffer (225 mM Mannitol, 75 mM Sucrose, 30 mM Tris [pH 7.4]). Cells were homogenized with a Dounce homogenizer on ice. Every 150 passes of the homogenizer, cells were centrifuged for 5 minutes at 600 x *g*, to remove non-lysed cells and nuclei. The integrity of the homogenized cells was examined under a light microscope until 80-90 % cell lysis was attained. The supernatant was collected as the post-nuclear supernatant (PNS) fraction. The clarified lysate was centrifuged at 7000 x *g* for 10 min at 4 °C, isolating the supernatant including ER-microsomes. The supernatant was collected, centrifuged at 20,000 x *g* for 30 mins at 4 °C to clear lysosomal and plasma membrane fractions. The resulting supernatant was further centrifuged at high speed, 100,000 x *g* for 1 hour in a swinging-bucket rotor SW-41 and Beckman Coulter ultracentrifuge. This results in the isolation of pure ER-microsomes (pellet) and cytosolic fraction (supernatant). PNS, cytosolic, and ER-microsomes were solubilized using 0.5 % (v/v) SDS and protein concentrations were determined by DC protein assay. An equivalent amount of ER-microsomes were used in the protease protection assay. Of four aliquots, one was left untreated, one was incubated with 1 % (v/v) Triton X-100, and the other two were incubated with 10 µg/mL proteinase K with or without 1 % (v/v) Triton X-100. All samples are incubated at 37 °C for 10 minutes before reaction termination with 2 mM phenylmethylsulfonyl fluoride (PMSF). 5X-Laemmli sample buffer was added to each sample before SDS-PAGE and immunoblot.

### Puromycilation translation assay

Protein synthesis was measured in HFT wildtype and UBXN1 KO cells utilizing the non-radioactive SUnSET assay^74^. HFT wildtype or UBXN1 KO cells were seeded onto a 6-well plate at 400 k cells /well to achieve 70-80 % confluence at the time of puromycin addition. Cells were incubated with 1 µM puromycin with or without 100 µg/ml cycloheximide for 30 minutes in complete medium. Samples treated with DTT or thapsigargin were pre-incubated with 1.5 mM DTT for 1 hour or 1 µM thapsigargin for 30 minutes before co-treatment of 1 µM puromycin with 1.5 mM DTT for 30 minutes in complete medium. At the end of the incubation period, cells were washed with ice-cold PBS and the plate was immediately placed on ice. Ice-cold RIPA buffer was added to the wells and cells were gently scraped into the buffer using a cell scraper. Lysates were placed on ice and vortexed for 10 seconds every 5 minutes for 30 minutes before centrifugation at 12,000 rpm for 10 minutes at 4 °C. Protein concentration was estimated using the DC assay (Bio-Rad). An equivalent amount of protein for each sample were resolved by SDS-PAGE and probed with the anti-puromycin antibody, clone 12D10.

### Polysome profiling

HEK-293T wildtype and UBXN1 KO cells were plated in a 150 mm dish and allowed to grow for two days before polysome isolation. For p97 depletion experiments, HEK-293T cells were plated in a 150 mm dish and forward transfected next day with siRNA against p97 or an siControl. 24 hours later the plate was split into two 150 mm dishes and the cells were grown for two additional days. The day before profiling, sucrose gradients were prepared with fresh 15 % and 60 % sucrose solutions using 1X low-salt buffer (5 mM Tris-HCl [pH 7.5], 2.5 mM MgCl_2_, 1.5 mM KCl, 1 mM PMSF). To generate a continuous gradient, 5 mL of 60 % sucrose solution was placed at the bottom of an ultra-clear Beckman Coulter centrifuge tube. 5 mL of 15 % sucrose solution was layered on top and the gradient was inverted horizontally and left on its side at room temperature for 3 hours. After 3 hours, the tube was brought back to its vertical position and stored at 4 °C overnight. When cells reached about 70-80 % confluency, cells were either pre-treated with 2 mM DTT for 60 minutes or left untreated. After treatment, cells were incubated with 100 ug/mL cycloheximide at 37 °C for 15-20 minutes. Cells were then washed twice with ice cold PBS supplemented with 100 ug/mL cycloheximide and 1 mM PMSF, scraped off the plate, and pelleted at 200 x *g* for 5 minutes at 4 °C. Cells were washed once with ice cold PBS, pelleted, and resuspended in 480 uL 1X low salt buffer supplemented with 100 ug/mL cycloheximide, 2 uM DTT, 1 mM PMSF, and 1 mM HALT protease inhibitor cocktail. 60 units of RNaseOUT Recombinant RNase Inhibitor (Invitrogen, 10777019) was added to each sample and vortexed for 5 seconds. Triton X-100 and sodium deoxycholate was added to each sample to a final concentration of 0.5 % and samples were vortexed for an additional 5 seconds. Samples were kept on ice for 5 minutes before centrifugation at 16,000 x *g* for 7 minutes at 4 °C. Samples were quantified at 260 nm in UV compatible plates on a multiwell plate reader. 20 units of each sample was loaded onto the continuous sucrose gradient and spun at 24,200 rpm for 3 hours 30 minutes using an SW-41 swing bucket rotor in a Beckman Coulter ultracentrifuge. After the spin, the samples were run on a continuous UA-6 detector, density gradient fractionator system with peak detection software. Values determined by the fraction collector were graphed on GraphPad Prism.

### TMT-based proteomics

#### Sample preparation, digestion, and tandem-mass tag (TMT) labeling

The TMT-based proteomics was performed exactly as previously described^85^. Briefly, 100 μg protein of each sample was obtained by cell lysis in lysis buffer (8 M Urea, 200 mM N-(2-Hydroxyethyl)piperazine-N′-(3-propanesulfonic acid) (EPPS) pH 8.5) followed by reduction using 5 mM tris(2-carboxyethyl)phosphine (TCEP), alkylation with 14 mM iodoacetamide and quenched using 5 mM dithiothreitol. The reduced and alkylated protein was precipitated using methanol and chloroform. The protein mixture was digested with LysC (Wako) overnight followed by Trypsin (Pierce) digestion for 6 hours at 37°C. The trypsin was inactivated with 30% (v/v) acetonitrile. The digested peptides were labelled with 0.2 mg per reaction of 4-plex TMT reagents (ThermoFisher scientific) (126, 127N, 129N, and 129C) at room temperature for 1 hour. The reaction was quenched using 0.5% (v/v) Hydroxylamine for 15 min. A 2.5 μL aliquot from the labeling reaction was tested for labeling efficiency. TMT-labeled peptides from each sample were pooled together at a 1:1 ratio. The pooled peptide mix was dried under vacuum and resuspended in 5% formic acid for 15 min. The resuspended peptide sample was further purified using C18 solid-phase extraction (SPE) (Sep-Pak, Waters).

#### Off-line basic pH reverse-phase (BPRP) fractionation

We fractionated the pooled, labeled peptide sample using BPRP HPLC^86^. We used an Agilent 1200 pump equipped with a degasser and a detector (set at 220 and 280 nm wavelength). Peptides were subjected to a 60-min linear gradient from 5% to 35% acetonitrile in 10 mM ammonium bicarbonate pH 8 at a flow rate of 0.25 mL/min over an Agilent 300Extend C18 column (3.5 μm particles, 2.1 mm ID and 250 mm in length). The peptide mixture was fractionated into a total of 96 fractions, which were consolidated into 24 super-fractions^87^. Samples were subsequently acidified with 1% formic acid and vacuum centrifuged to near dryness. Each consolidated fraction was desalted via StageTip, dried again via vacuum centrifugation, and reconstituted in 5% acetonitrile, 5% formic acid for LC-MS/MS processing.

#### Liquid chromatography and tandem mass spectrometry

Mass spectrometric data were collected on an Orbitrap Fusion mass spectrometer coupled to a Proxeon NanoLC-1000 UHPLC. The 100 µm capillary column was packed with 35 cm of Accucore 150 resin (2.6 μm, 150Å; ThermoFisher Scientific). The scan sequence began with an MS1 spectrum (Orbitrap analysis, resolution 120,000, 350−1400 Th, automatic gain control (AGC) target 5 x10^5^, maximum injection time 100 ms). Data were acquired for 150 minutes per fraction. SPS-MS3 analysis was used to reduce ion interference^88, 89^. MS2 analysis consisted of collision-induced dissociation (CID), quadrupole ion trap analysis, automatic gain control (AGC) 1 x10^4^, NCE (normalized collision energy) 35, q-value 0.25, maximum injection time 60 ms), isolation window at 0.7 Th. Following acquisition of each MS2 spectrum, we collected an MS3 spectrum in which multiple MS2 fragment ions were captured in the MS3 precursor population using isolation waveforms with multiple frequency notches. MS3 precursors were fragmented by HCD and analyzed using the Orbitrap (NCE 65, AGC 2.0 x10^5^, isolation window 1.3 Th, maximum injection time 150 ms, resolution was 50,000).

#### Data analysis

Spectra were converted to mzXML via MSconvert^90^. Database searching included all entries from the Human UniProt Database (downloaded: April 2016). The database was concatenated with one composed of all protein sequences for that database in the reversed order. Searches were performed using a 50-ppm precursor ion tolerance for total protein level profiling. The product ion tolerance was set to 0.9 Da. These wide mass tolerance windows were chosen to maximize sensitivity in conjunction with Comet searches and linear discriminant analysis^91, 92^. TMT tags on lysine residues and peptide N-termini (+229.163 Da for TMT) and carbamidomethylation of cysteine residues (+57.021 Da) were set as static modifications, while oxidation of methionine residues (+15.995 Da) was set as a variable modification. Peptide-spectrum matches (PSMs) were adjusted to a 1% false discovery rate (FDR)^93, 94^. PSM filtering was performed using a linear discriminant analysis, as described previously^92^ and then assembled further to a final protein-level FDR of 1% ^94^. Proteins were quantified by summing reporter ion counts across all matching PSMs, also as described previously^95^. Reporter ion intensities were adjusted to correct for the isotopic impurities of the different TMT reagents according to manufacturer specifications. The signal-to-noise (S/N) measurements of peptides assigned to each protein were summed and these values were normalized so that the sum of the signal for all proteins in each channel was equivalent to account for equal protein loading. Finally, each protein abundance measurement was scaled, such that the summed signal-to-noise for that protein across all channels equaled 100, thereby generating a relative abundance (RA) measurement.

Downstream data analyses for TMT datasets were carried out using the R statistical package (v4.0.3) and Bioconductor (v3.12; BiocManager 1.30.10). TMT channel intensities were quantile normalized and then the data were log-transformed. The log transformed data were analyzed with limma-based R package where p-values were FDR adjusted using an empirical Bayesian statistical. Differentially expressed proteins were determined using a log2 (fold change (WT *vs* UBXN1 KO)) threshold of > +/- 0.65.

#### Gene ontology (GO) functional enrichment analyses of proteomics data

The differentially expressed proteins were further annotated and GO functional enrichment analysis was performed using Metascape online tool (http://metascape.org)^96^. The GO cluster network and protein-protein interaction network generated by metascape and the STRING database (https://string-db.org/), respectively, were imported into Cytoscape software (v3.8.2) to add required attributes (fold changes, p-values, gene number, and conditions) and prepared for the visualization. Other proteomic data visualizations were performed using the RStudio software (v1.4.1103), including hrbrthemes (v0.8.0), viridis (v0.6.1), dplyr (v.1.0.7), and ggplot2 (v 3.3.5).

#### Quantification and statistical analysis

Immunoblots were quantified using densitometry on FIJI and protein levels were normalized to the loading control for that experiment (GAPDH or *β*-actin). Fold changes, SEM, and statistical significance was calculated using GraphPad Prism (version 9.4). Statistical tests performed are indicated in figure legends and include unpaired two-tailed Student’s *t*-test or one-way ANOVA with post-hoc analysis indicated in figure legends.

## Acknowledgements

We thank Peter Juo and Michelle Johnson and Rakesh Ganji for critical reading of the manuscript. We are grateful to Peter Walter (University of California San Francisco and Altos Labs) for the GFP-IRE1 construct, and Mals Mariappan (Yale University) for the FLAG-AGAL construct. This work is supported by the NIH grants GM127557, NS123631 and a Research Scholar Grant RSG-19-022-01 from the American Cancer Society to M.R. This work was funded in part by NIH/NIGMS grant GM67945 (S.P.G.) and R01 GM132129 (J.A.P.).

## Respective Contributions

B.A and M.R conceived the studies. R.G performed TMT proteomics and data analysis, S.M performed aggresome studies. J.P and S.P.G assisted with proteomic studies. B.A and M.R wrote the manuscript.

## Request for reagents

Please contact the corresponding author, M.R for reagent requests.

## Data availability

All raw proteomic data will be made available on public servers upon acceptance of the manuscript. Any other data is available from the corresponding author upon request.

